# Frequent intertrophic transmission of *Wolbachia* by parasitism but not predation

**DOI:** 10.1101/2024.03.20.585917

**Authors:** Zhi-Chao Yan, Lan-Da Qi, Han-Le Ji, Xiao-Xiang Wang, Xiao-Yue Hong, Yuan-Xi Li

## Abstract

*Wolbachia* is one of the most pervasive symbionts, estimated to infect ∼50% of arthropod species. It is primarily transmitted vertically, inducing a variety of fascinating reproductive manipulations to promote its spread within host populations. However, incongruences between host and *Wolbachia* phylogenies indicate substantial horizontal transmissions, the mechanisms of which are largely unexplored. By systematically analyzing *Wolbachia* surface protein (*wsp*) sequences on NCBI, we found that parasitism, not predation, is the primary route of intertrophic *Wolbachia* transmission. This conclusion held after accounting for sampling bias. One example of frequent *Wolbachia* transfers is between egg parasitoid wasps, *Trichogramma*, and their lepidopteran hosts. Moreover, both bioinformatics and experimental results showed that *Wolbachia* from the parasitoid wasp *Encarsia formosa* can be transmitted to its whitefly host *Bemisia tabaci*, through unsuccessful parasitism. Once *En. formosa Wolbachia* is transferred to whiteflies, it can be vertically transmitted within whiteflies and induce fitness costs. To our knowledge, this is the first compelling evidence that *Wolbachia* can be transmitted from parasitoid wasps to their hosts, revealing the bidirectional nature of *Wolbachia* transfers between parasitoids and their hosts. Overall, our findings enrich the current understanding of the horizontal transmission of *Wolbachia* and shed new light on its ecology and evolution.

## Introduction

Symbiosis with microbes, ranging from parasitism to mutualism, is prevalent in both plants and animals (*1, 2*). The ubiquity of microbial symbionts is likely attributed to their profound impact on host biology, including host survival, development, immunity, reproduction, and even behavior (*2, 3*). The transmission mode of symbionts is a key factor in shaping the ecology and evolution of both symbionts and their hosts (*4–6*). In addition to vertical transmission, incongruence between symbionts and host phylogenies indicates a large number of horizontal symbiont transfers across species (*1*). These events are important, as they allow symbionts to expand their host range and enable hosts to acquire new symbionts and alter their fitness.

*Wolbachia* (Rickettsiales, Alphaproteobacteria) is intracellular gram-negative bacteria and one of the most famous endosymbionts that infests ∼50% arthropod and several filarial nematode species (*7–9*). On the one hand, *Wolbachia* can induce a range of fascinating phenotypes, including a variety of reproductive manipulations, provision of nutrients, and alteration of host behavior, thus facilitating its spread among populations (*7*). On the other hand, although *Wolbachia* is primarily maternally transmitted, there are widespread and frequent horizontal transfers across hosts (*10, 11*). Together, these characteristics make *Wolbachia* likely the most common microbe on Earth in terms of the number of species it infects (*7, 8*).

*Wolbachia* has also received much attention for its applications in controlling pests and vector-borne diseases (*8*). Specific strains of *Wolbachia* are artificially transfected into target pests and subsequently released into the field to either suppress pest populations or replace populations to depress the spread of vector-borne diseases (*12–15*). Understanding how *Wolbachia* spreads horizontally is critical in assessing its successful application and potential risks (*16*). This is because the released *Wolbachia* may leak into natural pest populations, frustrating population suppression strategies based on the cytoplasmic incompatibility (CI) of *Wolbachia*. It may also spread to nontarget organisms, potentially disrupting their population dynamics. Therefore, the lack of a thorough understanding of *Wolbachia* transmission and its consequences could hinder its broader application (*16*).

Despite the extensive interest in the horizontal transmission of *Wolbachia*, our understanding of this subject remains incomplete (*17*). Similar to other symbionts, *Wolbachia* host shifts may occur through three main routes: parasitism, predation, and shared plant or other food sources (*17*). However, it is important to note that these are not the only routes through which transmission may occur, and the specific contributions of each to the overall process of host shift are not yet fully understood. Multiple surveys report a significant similarity in *Wolbachia* sequences between parasitoid wasps and their respective hosts, suggesting that parasitism may serve as a primary route for *Wolbachia*’s host shift (*18–23*). However, there is still a lack of systematic statistical analyses to support this hypothesis. For the intertrophic transmission of *Wolbachia* between parasitoid wasps and their hosts, experimental evidence has shown that parasitoid wasps can acquire hosts’ *Wolbachia* and vertically transmit them for several generations (*24*). However, it remains uncertain whether *Wolbachia* can be transferred from parasitoid wasps to their hosts. Some have argued that the transfer of *Wolbachia* between parasitoid wasps and their hosts is unidirectional, from host to wasp, as all parasitoid wasps emerge from their hosts, but parasitized hosts are eventually killed or castrated if not killed immediately (*19, 25, 26*). However, it is common in nature for hosts to survive parasitoid attacks (*27–29*). For example, whiteflies can survive after attacks of *Eretmocerus* parasitoids (*27*). These parasitoids can serve as phoretic vectors for the transmission of *Wolbachia* among whitefly populations, with contamination of *Wolbachia* on their mouthparts and ovipositors during the probing process (*27*).

In this study, our objective was to elucidate the role of parasitism and other potential routes in *Wolbachia* horizontal transmission and to investigate whether *Wolbachia* can be transferred from parasitoids to their hosts. We conducted a systematic survey of *Wolbachia* surface protein (*wsp*) sequences from the NCBI database and executed experiments using the whitefly *Bemisia tabaci* and its parasitoid wasp *Encarsia formosa*. Our study illuminates the crucial role of parasitism in *Wolbachia* intertrophic transmission, demonstrates the bidirectional nature of *Wolbachia* transfer between parasitoids and their hosts, and thus expands the current understanding of *Wolbachia* horizontal transmission.

## Results

### *Wolbachia* is frequently transmitted between parasitoid wasps and their hosts

To investigate potential horizontal transmission of *Wolbachia*, we retrieved 4685 *wsp* sequences from the NCBI database, and species interaction relationships were extracted from the GloBI database (for details, see Methods and Materials). Out of these 4685 sequences, 4253 could be assigned to 1377 species. We constructed a phylogenetic tree of *wsp* sequences (Fig. 1a) and extracted the minimum genetic distances of *wsp* between every species pair. Based on the relationships between species, we defined the species pairs into categories “Parasitism”, “Plant-sharing”, “Predation” and “Others”. Among these species pairs, 16.5% (53 out of 321) in “Parasitism” pairs had the minimum interspecific *wsp* distances less than 0.01 (i.e., > 99% identity). This proportion is significantly greater than the 6.3% (145 out of 2294) in “Plant-sharing” pairs, the 0.9% (13 out of 1446) in “Predation” pairs, and the 1.5% (14120 out of 943315) in “Others” pairs (*χ^2^* test; all comparisons: *p* < 1e^-5^). Consistently, the minimum interspecific *wsp* distances in “Parasitism” relationships were significantly shorter than those in “Plant-sharing”, “Predation”, and “Other” relationships (Fig. 1b; Mann Whitney U test (MWUT); all comparisons: *p* < 1e-12). The minimum *wsp* distances in “Plant-sharing” were also significantly smaller than those in “Others” (Fig. 1b; MWUT; *W* = 730215564, *p* < 2.2e^-16^). However, the minimum *wsp* distances showed no significant difference between “Predation” and “Others” (Fig. 1b; MWUT; *W* = 699285646, *p* = 0.096).

**Fig. 1.**
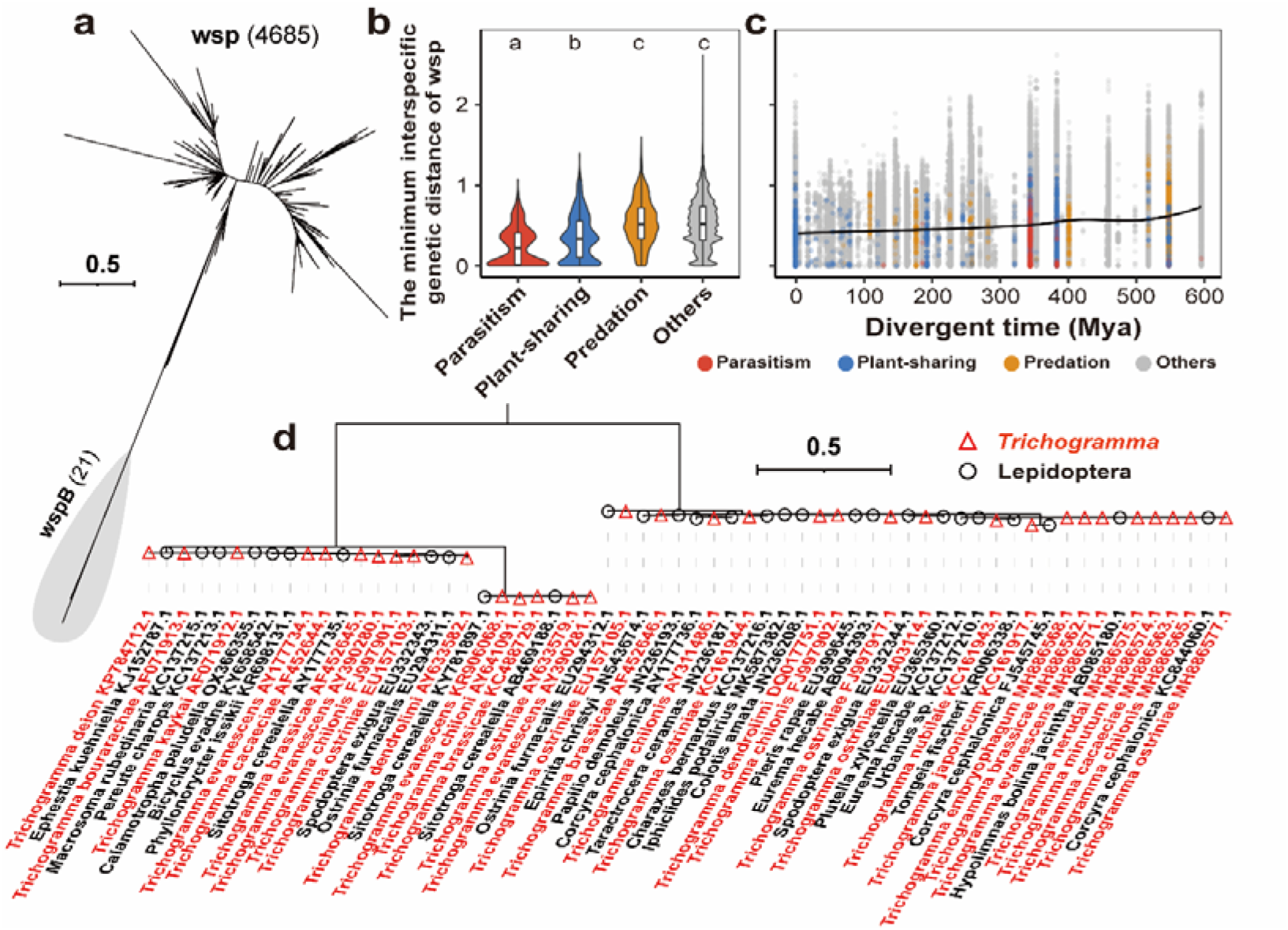
Parasitism facilitates interspecific horizontal transfer of *Wolbachia*. **(a)** Phylogeny of *Wolbachia* surface protein (*wsp*) genes retrieved from NCBI. **(b)** Effect of parasitism, plant-sharing, and predation on the genetic distance of *wsp* between species. **(c)** Relationship between species divergence time and the genetic distance of *wsp* between species. **(d)** Phylogeny of representative *wsp* sequences from *Trichogramma* wasps and lepidopterans. Color red and black represent *wsp* sequences from *Trichogramma* wasp and lepidopterans, respectively.

To test whether these effects were merely due to sampling bias, we further obtained divergent times from TimeTree for 95.2% (901,615 out of 947,376) of the species pairs. As expected, the minimum interspecific *wsp* distance increased as the divergent time increased (Fig. 1c; Spearman’s correlation; *ρ* = 0.14, *p* < 2.2e^-16^). Considering the impact of divergent time, linear regression analyses were conducted with the *wsp* distance as the dependent variable and divergent time as the independent variable. We found that both “Parasitism” and “Plant-sharing” had significant effects (both tests: *p* < 2e^-16^). The estimated effect of “Parasitism” (-0.28) was more profound than that of “Plant-sharing” (-0.13) (*p* = 3.9e^-16^). Similar results were observed even after considering phylogenetic correction (Supplementary Note1). Given that we cannot obtain divergence times for all species pairs, especially for those closely related, we also classified the species pairs according to their last common ancestor. We divided all species pairs into six categories based on whether the two species belonged to the same genus, family, order, class, phylum, or kingdom. Similar results were observed (Supplementary Note 2). To further rule out potential influences of sampling bias, subsampling analyses were conducted using three methods: 1) shuffling species relationships, 2) subsampling by controlling divergent time, and 3) subsampling by controlling the last common ancestor (for details, see Methods and Materials). For all three methods, both “Parasitism” and “Plant-sharing”, but not “Predation”, exhibited significantly shorter minimum interspecific distances of *wsp* than randomly generated controls (Fig. S1; both “Parasitism” and “Plant-sharing”: *p* < 0.001; “Predation”: *p* > 0.05). These results confirmed that parasitism and plant-sharing promote interspecific transfers of *Wolbachia*, rather than being due to sampling bias.

An example of frequent *Wolbachia* transfers between parasitoids and their hosts is observed in *Trichogramma* wasps and their lepidopteran hosts (Fig. 1d). *Trichogramma* is a genus of generalist egg parasitoids, targeting mainly lepidopterans such as moths and butterflies (*30*). Fig. 1d displays representative *wsp* sequences from *Trichogramma* and lepidopterans, illustrating potential interspecific transitions of *Wolbachia*. In the analyzed dataset, 16 out of 23 (69.6%) *Trichogramma* species exhibited the minimum interspecific *wsp* distance of less than 0.01 with at least one lepidopteran species. Similarly, 79 out of 254 (31.1%) surveyed lepidopteran species displayed a minimum interspecific *wsp* distance of less than 0.01 with at least one *Trichogramma* species. These results suggest frequent *Wolbachia* transitions between *Trichogramma* wasps and lepidopterans.

Collectively, these findings support that *Wolbachia* are frequently transmitted between parasitoid wasps and their hosts.

### *Wsp* phylogeny suggests transfer directions of *Wolbachia* between the whitefly *B. tabaci* and its parasitoid wasps

However, the interspecific identities of *wsp* between parasitoids and their hosts typically offer no clues about the direction of *Wolbachia* transmission. We noticed frequent *Wolbachia* transfers between the whitefly *B. tabaci* and its parasitoid wasps (Fig. 2a). Many of the *wsp* sequences from *B. tabaci* and its parasitoids shown in Fig. 2a were generated in our previous field study (*31*). These previously published sequences were reanalyzed as part of the dataset retrieved from NCBI for the present study. Notably, one of the juvenile parasitoids of *B. tabaci* is *En. formosa*. *Wolbachia* induces parthenogenesis and has evolved into an obligate symbiont in *En. formosa*, exhibiting a 100% infection rate (*32, 33*). This system presents a unique opportunity to infer the directions of *Wolbachia* transmission between parasitoids and their hosts. One clade of the *wsp* phylogeny contained 99 sequences from *B. tabaci*, 14 sequences from *Encarsia* and *Eretmocerus* parasitoid wasps of *B. tabaci*, and 8 other sequences (Fig. 2a). Based on ancestral state reconstruction, the ancestral state of the Order for these *wsp* sequences was Hemiptera with a high likelihood (100.00%), while all other orders were negligible (<0.001%). This overwhelming probability strongly suggests that the transmission direction of *Wolbachia* in this clade is predominantly from *B. tabaci* to its parasitoid wasps. Notably, in another clade of *wsp*, nine *En. formosa* and five *B. tabaci wsp* sequences are clustered together, along with one *wsp* sequence detected in cotton and 12 sequences from other parasitoid wasps of *B. tabaci*, such as *En. inaron*, *En. lutea*, *En. bimaculata*, *Er. mundus*, and *Amitus hesperidum* (Fig. 2a). The ancestral state of the Order for these *wsp* sequences was estimated to be Hymenoptera with a near certain likelihood of 100.00%. Given that *Wolbachia* is obligate with a 100% infection rate in *En. formosa*, it is reasonable to infer that the transmission direction of *Wolbachia* in this clade was from *En. formosa* to *B. tabaci*.

**Fig. 2.**
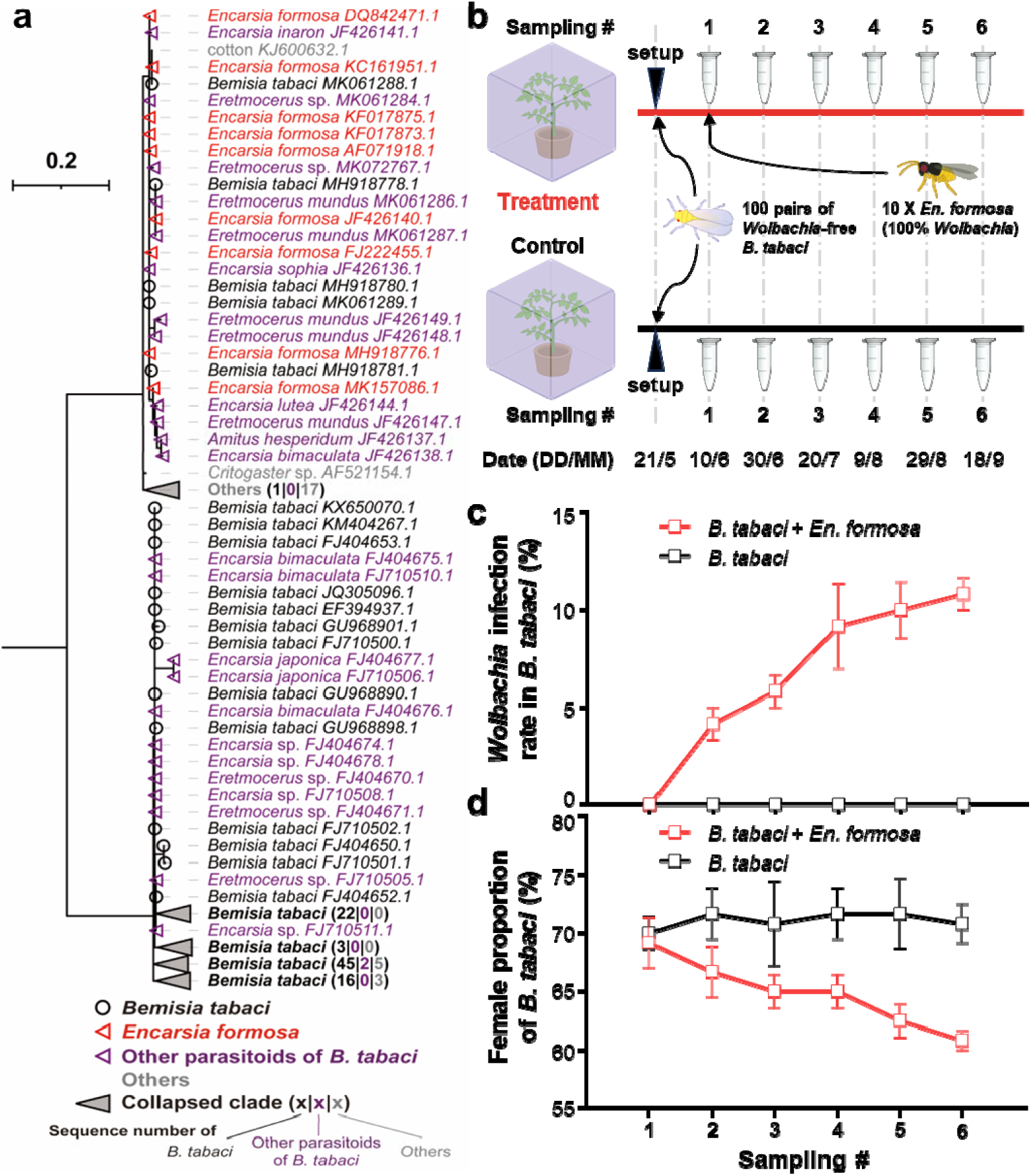
Transfer of *Wolbachia* from the parasitoid wasp, *Encarsia formosa*, to its host, *Bemisia tabaci*. **(a)** Phylogenetic analysis of the *wsp* gene from *B. tabaci* and its parasitoid wasps. The colors black, red, purple, and gray represent *wsp* sequences from *B. tabaci*, *En. formosa*, other parasitoids of *B. tabaci*, and other species, respectively. Previously published *wsp* sequences from Qi et al. (*31*) are included in the dataset and reanalyzed here. **(b)** Scheme of the experimental design for studying the transmission of *Wolbachia* from *En. formosa* to *B. tabaci*. **(c–d)** Effects of *En. formosa* on **(c)** the infection rate of *Wolbachia* and **(d)** the proportion of females in *B. tabaci* populations. Data are presented as the means ± standard errors (SE) (n = 3).

### *Wolbachia* can be transmitted from the parasitoid wasp *En. formosa* to its whitefly host in cage experiments

*Bemisia tabaci* is a species complex of at least 40 cryptic species (*34*). Infection rates of *Wolbachia* reported across multiple populations within these cryptic species exhibit dramatic fluctuations, ranging from 0% to 100% (*34–37*). To investigate whether *Wolbachia* can be transmitted from parasitoid wasps to their whitefly hosts, we first established a *Wolbachia*-free iso-female line of *B. tabaci* Q biotype. Subsequently, we conducted outdoor cage experiments using the *Wolbachia*-free *B. tabaci* and its parasitoid *En. formosa*, which was 100% infected with *Wolbachia* (Fig. 2b). Notably, each cage setup was replicated three times to ensure experimental rigor. After introducing *En. formosa*, the *Wolbachia* infection rate in whiteflies increased from zero to 4.17% after 20 days and reached 10.83% after 100 days (Fig. 2c; ANOVA; *F*_5,12_=11.44, *p* < 0.001). Correspondingly, the female ratio of whiteflies decreased from 69.17% to 60.83% (Fig. 2d; ANOVA; *F*_5,12_=3.14, *p* = 0.048). In contrast, the whitefly population with no exposure to wasps maintained a zero infection rate of *Wolbachia* (Fig. 2c), and the female ratio showed no significant changes (Fig. 2d; ANOVA; *F*_5,12_ = 0.076, *p* = 0.99). Additionally, the Sanger sequencing results showed that *wsp* sequences from both whitefly and *En. formosa* were identical (Fig. S2). These results indicate that *Wolbachia* from *En. formosa* can be rapidly transmitted to its whitefly host.

### Parasitism failure transmits *Wolbachia* from the parasitoid wasp *En. formosa* to its whitefly host

We hypothesized that *Wolbachia* was transmitted from parasitoid wasps to their hosts through unsuccessful parasitism. To test this hypothesis, we performed parasitism experiments on wasp individuals and applied irradiation treatment to wasps to reduce their parasitism success rate. After 60 Gy irradiation, the fecundity of *En. formosa* in 12 h significantly decreased (Fig. 3a; Student’s t test; *t* = 12.91, *p* < 0.001), and the parasitism success rate on whiteflies drastically declined from 78.8% to 9.9% (Fig. 3b; Student’s t test; *t* = 13.09, *p* < 0.001). Correspondingly, the *Wolbachia* infection rate increased from 7.5% to 78.3% in whiteflies that survived from parasitism (Fig. 3c; Student’s t test; *t* = 6.61, *p* < 0.001), despite a decrease in the *Wolbachia* titer in *En. formosa* post irradiation (Fig. S3). More details can be found for irradiation treatments at different dosages (Supplementary Note 3). Fluorescence in situ hybridization (FISH) assays further showed that *Wolbachia* was injected into the host nymphs along with *En. formosa* eggs (Fig. 3d–f and S4). These results indicate that parasitism failure can transfer *Wolbachia* from the parasitoid wasps *En. formosa* to their whitefly hosts.

**Fig. 3.**
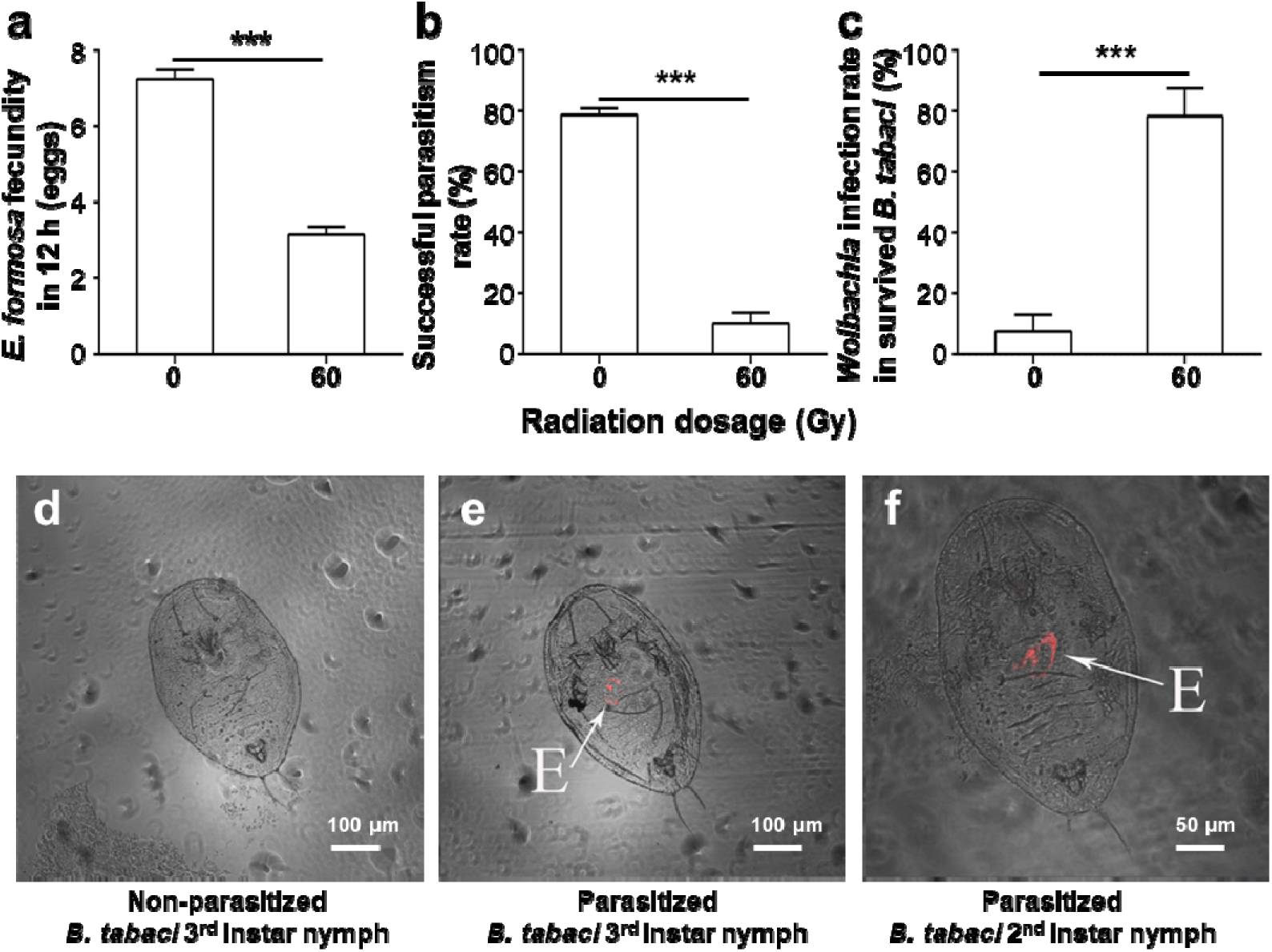
Parasitism failure mediates *Wolbachia* transfer from *En. formosa* to *B. tabaci*. **(a–c)** Effects of 60 Gy radiation on **(a)** the fecundity of *En. formosa* over a 12-hour period, **(b)** the rate of successful parasitism, and **(c)** the *Wolbachia* infection rate in surviving whiteflies. Data are presented as the means + SEs (n = 20). ***: *p* < 0.001. **(d–f)** Fluorescence in situ hybridization (FISH) visualization of *Wolbachia* in nonparasitized and parasitized 3^rd^ instar nymphs, as well as parasitized 2nd instar nymphs of *B. tabaci*. The images present a combination of bright field and fluorescence. E: injected eggs from *En. formosa*.

### Vertical transmission and fitness cost of *Wolbachia* in whiteflies after horizontal transfer from *En. formosa*

Next, we investigated the vertical transmission and fitness cost of *Wolbachia* in its new host, *B. tabaci*, following its horizontal transfer from *En. formosa*. The vertical transmission rate varied from 22.2% to 33.3% across generations in whiteflies, with a slight but statistically insignificant increase from G_1_ to G_5_ (Fig. 4b; ANOVA; *F*_4,40_ = 2.09, *p* = 0.10). Moreover, *Wolbachia*-targeted FISH signals were detected in G3 whiteflies. In nymphs, the signals were observed in structures morphologically resembling bacteriocytes (Fig. 4c and S5). In adults, they were detected in the abdomen of males and the ovaries of females (Fig. 4d, e and S5). *Wolbachia*-free whiteflies that had not been parasitized by *En. formosa* served as negative controls. These whiteflies were processed in parallel with the *Wolbachia*-positive samples and imaged using identical exposure settings. No *Wolbachia*-specific FISH signals were detected in the negative controls (Fig. S6). MLST typing confirmed that the *Wolbachia* strain introduced into the whiteflies was identical to the strain from *En. formosa* (Fig. S7).

**Fig. 4.**
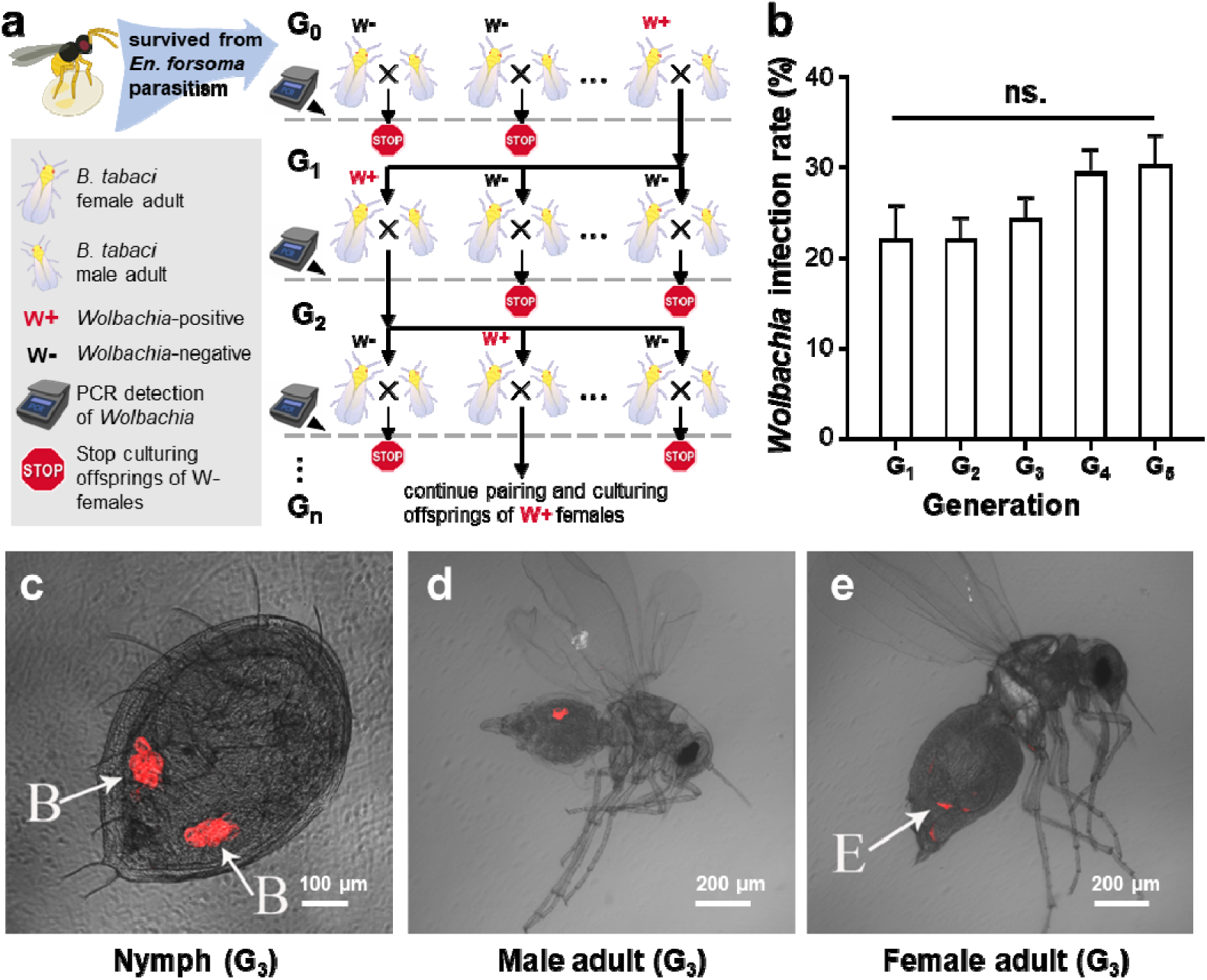
Vertical transmission of *Wolbachia* from *En. formosa* in *B. tabaci*. **(a)** Scheme of the study design for *Wolbachia* vertical transmission in *B. tabaci*. Adult whiteflies that survived from *En. formosa* parasitism were denoted as G_0_. After pairing and oviposition, the infection status of *Wolbachia* in the female parent was examined. Only the offsprings from *Wolbachia*-infected female whiteflies were maintained. **(b)** The vertical transmission rate of *Wolbachia* across five generations in *B. tabaci*. Data are presented as the means + SEs (n = 9). ns.: no significant differences. **(c–e)** FISH visualization of *Wolbachia* in G_3_ *B. tabaci* **(c)** nymph, **(d)** male adult, and **(e)** female adult. The images present a combination of bright field and fluorescence. B: structures morphologically resembling bacteriocytes; E: eggs in the ovary of a female whitefly.

We then examined the impact of the introduced *Wolbachia* on the fitness of whiteflies. Compared to uninfected females, *Wolbachia*-infected females showed decreased fecundity (Fig. 5a; Student’s t test; *t* = 8.51, *p* < 0.001). Moreover, offspring from infected mothers exhibited a diminished egg hatching rate (Fig. 5b; Student’s t test; *t* = 8.33, *p* < 0.001), a lower survival rate among nymphs (Fig. 5c; Student’s t test; *t* = 13.54, *p* < 0.001), and a decreased ratio of females in adults (Fig. 5d; Student’s t test; *t* = 12.29, *p* < 0.001), although there was no significant difference in the development time from egg to adult (Fig. 5e; Student’s t test; *t* = 1.51, *p* = 0.16). We also investigated the effects of paternal infection with *Wolbachia*. Regardless of whether the female was infected with *Wolbachia*, the infection status in males showed no significant influence on their fecundity, the hatching rate of their offspring’s eggs, the survival rate of their nymph offspring, or the female ratio of their adult offspring (Table 1; Student’s t test; all comparisons: *p* > 0.05).

**Fig. 5.**
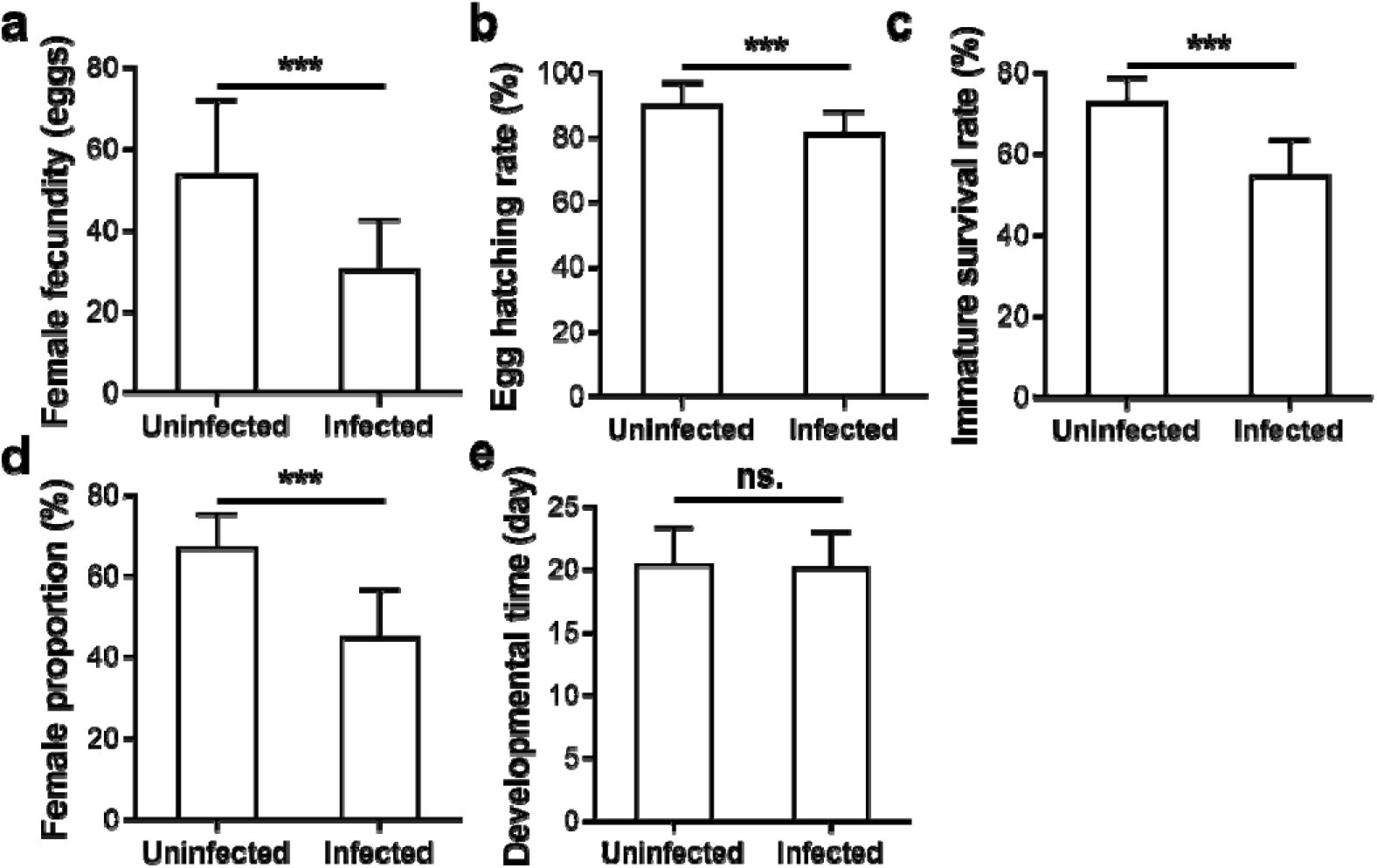
Fitness costs in *B. tabaci* induced by *Wolbachia* from *En. formosa*. **(a–e)** The effects of *En. formosa Wolbachia* on *B. tabaci* **(a)** female fecundity, **(b)** egg hatching rate, **(c)** immature survival rate, **(d)** female proportion, and **(e)** developmental time. Data are presented as the means + SDs (n = 60). ***: *p* < 0.001; ns.: no significant differences.

**Table 1.**
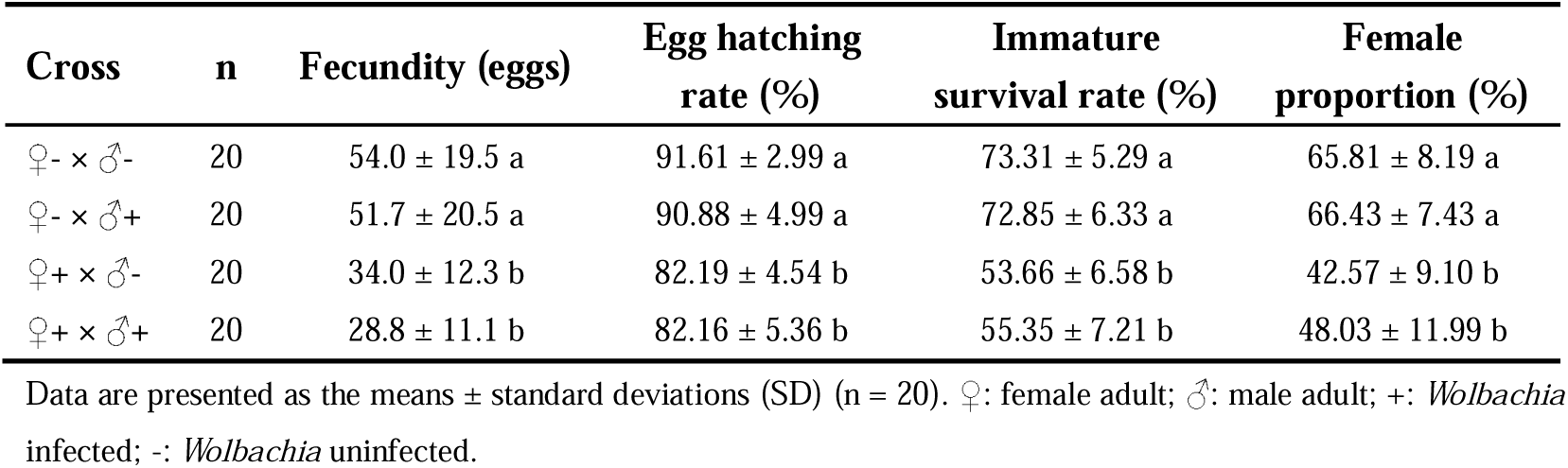
Fitness in *Bemisia tabaci* of different cross combinations.

Collectively, these results indicate that *Wolbachia* from *En. formosa*, when shifted into the new host *B. tabaci*, exhibits a low rate of vertical transmission and a substantial fitness cost, without apparent reproductive manipulation phenotypes.

## Discussion

### The effect of parasitism on *Wolbachia* horizontal transmission

As with previous studies, we utilized the sequence similarities of *wsp* to infer potential horizontal transfers of *Wolbachia*. Here, we systematically investigated all *wsp* sequences in the NCBI database, enabling us to examine the effect of potential factors such as parasitism, plant-sharing, and predation. Our findings clearly indicate that parasitism exhibits shorter interspecific *wsp* distances than plant-sharing and predation. For example, we found that some *wsp* sequences from multiple *Trichogramma* species and their various lepidopteran hosts are almost identical. A previous study has found common horizontal transmission between butterflies and moths, and speculated that parasitoids might be one of the routes that contribute to the horizontal transmission (*38*). Our results suggest potential horizontal transmission between *Trichogramma* wasps and their lepidopteran hosts, and *Trichogramma*, which is a kind of egg parasitoids of many lepidopterans, may act as vectors facilitating the transfer of *Wolbachia* among lepidopteran hosts.

Moreover, the fact that herbivores sharing plants have identical *wsp* sequences does not necessarily imply plant-mediated horizontal transfer of *Wolbachia*. This is because species that share the same plants often have recent divergent times (Fig. 1c), which increases the potential for hybridization and also sharing common parasitoid wasps. Therefore, the plant-sharing category overestimates the extent of plant-mediated *Wolbachia* horizontal transmission, further supporting the notion that parasitism is the primary route of *Wolbachia* horizontal transmission.

### Directions of *Wolbachia* transmission between parasitoids and their hosts

However, investigations based on *Wolbachia* sequence similarity have significant limitations. First, the PCR detection of *wsp* does not necessarily indicate horizontal transfer of *Wolbachia* (*26*). Instead, it could merely represent contamination that arises during predation or parasitism. The contamination can be *Wolbachia*-infected tissues or even just fragmented *Wolbachia* DNA, which could be found on the surface, within the gut, or inside the body cavity (as in the case of parasitized hosts).

Experimental confirmations of *Wolbachia* horizontal transfer remain relatively rare, with only a limited number of documented cases (*24, 27, 39, 40*). Additionally, some experiments have found no evidence of horizontal transmission of *Wolbachia* (*41–44*). For example, Chiel et al. detected no evidence of *Wolbachia* horizontal transmission within or between trophic levels in *B. tabaci* and its parasitoid *Eretmocerus* sp. nr. *emiratus* (*44*).

Second, determining the direction of transfer is challenging based solely on the identity of *Wolbachia* strains. In certain exceptional cases, inferences of transfer direction might be drawn from the prevalence of *Wolbachia* infection in the two species (*17*). For example, *Wolbachia* is obligate in *En. formosa* and exhibits a 100% infection rate across all populations (*33*). This unique instance of horizontal *Wolbachia* transfer between *En. formosa* and its whitefly hosts provides compelling evidence, suggesting that *Wolbachia* is likely transmitted from the parasitoid wasp to its host, rather than the reverse.

To verify *Wolbachia* transmission from parasitoid wasps to their hosts, we conducted outdoor cage experiments and indoor tests using *Wolbachia*-free whitefly *B. tabaci* and its parasitoid *En. formosa*. *Wolbachia* was detected by nested PCR in whitefly adults (G_0_) that survived parasitism. *Encarsia formosa* parasitizes *B. tabaci* nymphs, which undergo a pseudo-pupal stage to reach adulthood (*32, 45*). In our experiments, PCR detection of *Wolbachia* typically occurred 3–7 days post-parasitism, minimizing risks of contamination from parasitism. *Wolbachia* was also detected by PCR in subsequent generations (G1–G5) of whiteflies and induced notable fitness costs. PCR sequencing confirmed that the *Wolbachia* strain in the *B. tabaci* matched that of *En. formosa*. Furthermore, FISH assays revealed a tissue-specific distribution of *Wolbachia* in both nymphs and adults of the whiteflies, matching previously reported patterns (*46*). Collectively, these findings provide compelling evidence that *Wolbachia* from *En. formosa* can be horizontally transmitted to *B. tabaci*, beyond mere DNA contamination.

Although previous research has demonstrated that parasitoids can acquire *Wolbachia* from their hosts, the reverse direction of transmission, from parasitoid to host, has been largely overlooked and lacks supportive experimental evidence. One possible reason for this oversight is that all parasitoids emerge from their hosts, but hosts are eventually killed by the parasitism of parasitoid wasps (*19, 25, 26*). However, parasitoid wasps’ success in parasitizing their hosts does not always reach 100% (*28*). Some hosts can manage to survive after parasitism. Various factors can influence the outcome of parasitoid-host interactions, including environmental conditions, the species and genotype of both wasps and hosts, the host’s age, and the presence of symbiotic bacteria within the host (*29, 47, 48*). Moreover, we used radiation to reduce the parasitism success rate, which notably enhanced the transfer of *Wolbachia* from *En. formosa* to its whitefly host.

### Potential intertrophic transmission network of *Wolbachia*

A previous study reported that parasitoid wasps can act as vectors to transmit *Wolbachia*, without the necessity of being infected themselves (*27*). Through the probing actions of the *Eretmocerus* parasitoid, *Wolbachia* can be transmitted among whitefly hosts (*27*). This is often referred to as the ‘dirty needle’ model. Conversely, hosts can also serve as vectors for *Wolbachia* transmission among parasitoid wasps. *Wolbachia* can be transferred from infected to noninfected *Trichogramma* wasps through superparasitism (*39, 49*). However, the *Wolbachia* transmission of these two modes is restricted within the same trophic level. In contrast, the transmission of *Wolbachia* between parasitoid wasps and hosts can cross trophic levels. Our findings, when combined with existing knowledge, suggest that the intertrophic transmission of *Wolbachia* is bidirectional between parasitoid wasps and their hosts. This greatly enhances our understanding of the horizontal transmission of *Wolbachia*.

Interestingly, we found on NCBI that a strain of *Wolbachia* detected in the cotton plant was identical to the *Wolbachia* from *En. formosa*, based on the *wsp* sequence and MLST typing (Fig. 2 and S7). This *Wolbachia* strain was probably transmitted to the cotton plant from the feeding of whiteflies (*40*). Given that *En. formosa* is 100% infected with its obligate *Wolbachia* strain, a possible transmission route could be from *En. formosa* to whiteflies via parasitism and then to the cotton plant through the feeding of whiteflies. This finding indirectly supports our conclusion that *Wolbachia* can be transmitted from parasitoid wasps to their hosts. This suggests that once the *Wolbachia* of parasitoids is transmitted to herbivorous hosts, it may further spread to host plants. The reverse transmission route can also be possible. It is likely that *Wolbachia*’s widespread and complex horizontal transmission network is established through such bidirectional transmissions across multiple trophic levels, e.g., “plant-herbivore-parasitoid-hyperparasitoid”. Further investigations are needed to test these hypotheses.

### *Wolbachia* establishment after host transfers

Moreover, the physical transfer of *Wolbachia* often represents merely the first step of its establishment in a new host (*17*). Several subsequent steps are required for *Wolbachia* to establish itself within new species, e.g., entry into germ cells, vertical transmission, and mechanisms that promote its spread within the population (*17*). First, we found that *Wolbachia* from *En. formosa* was enriched in the ovaries of whiteflies and vertically transmitted after entering *B. tabaci*. However, the vertical transmission efficiency is low, ranging from 22.2% to 33.3%. We also noted an absence of reproductive manipulations by the newly introduced *Wolbachia* in whiteflies. The reduced female ratio after infection does not support the induction of parthenogenesis, feminization or male-killing by *Wolbachia*. We neither observed cytoplasmic incompatibility, where the mating of infected males and uninfected females resulted in reduced offspring hatching.

In contrast, the introduced *Wolbachia* from *En. formosa* reduced whiteflies’ egg laying and hatching, larval survival, and female proportion, demonstrating significant fitness costs in the new host. This is likely due to *Wolbachia*’s coevolution with its host in *En. formosa*, which may have led to the loss of its ability to colonize new host species. Given the low vertical transmission rate, high fitness cost, and lack of clear reproductive manipulations, it is reasonable to predict that the spread of *Wolbachia* in its new host population will be limited. Finally, these factors, together with the frequency of *Wolbachia* introductions by parasitoids and its spread via parasitoid or plant vectors, shape the dynamics and equilibrium of *Wolbachia*. These dynamics could shift with the emergence of reproductive manipulation or other beneficial phenotypes that promote *Wolbachia* spread, probably through gene mutation, recombination or horizontal gene transfer within *Wolbachia* (*2*). There are still many questions waiting to be further studied in these steps of *Wolbachia* host shifts.

## Conclusions

By investigating *wsp* sequences from the NCBI database, we found frequent intertrophic transmission of *Wolbachia* by parasitism but not predation. Combining bioinformatics and experimental approaches, we demonstrated that *Wolbachia* can be transmitted from the parasitoid wasp *En. formosa* to the host *B. tabaci*. To our knowledge, this is the first compelling evidence that *Wolbachia* can be transmitted from parasitoid wasps to their hosts, thus revealing the bidirectional nature of *Wolbachia* transfers between parasitoids and their hosts. These findings enrich our knowledge of the *Wolbachia* transmission network and have significant implications for understanding the ecology of *Wolbachia*, as well as for evaluating the release of *Wolbachia* in pest control.

## Materials and Methods

### *Wsp* sequence retrieval and phylogenetic analyses

TBLASTN was conducted against the NCBI “nr/nt” database using the *wsp* protein AAS14719.1 from *Wolbachia* of *Drosophila melanogaster* (July 2023). Default settings were used with a maximum target sequence of 5000 and the organism limitation of *Wolbachia* (taxid:953). The retrieved dataset includes *wsp* sequences generated by our laboratory from field collected samples reported previously (*31*) and deposited in GenBank, together with sequences submitted by other research groups. These previously published sequences (*31*) were re analyzed in the current study and were not newly generated for the horizontal transmission experiments described herein. The sequences were filtered to remove those shorter than 300 bp or containing premature stop codons. *Wsp* sequences were translated into proteins, aligned using MAFFT v7.475 (*50*), and reverse translated into codons using PAL2NAL v14 (*51*). The phylogenetic trees were constructed using IQ-Tree v2.2.0 (*52*), with the best model selected by the built-in ModelFinder (*53*). The phylogenetic trees were visualized using iTOL v6 (*54*).

### Statistics for genetic distances of *wsp*

Genetic distances of *wsp* sequences were extracted from the *wsp* phylogeny using a custom Python script. Species interaction relationships were extracted from the GloBI database (August 2023) (*55*). Parasitic associations were extracted using interaction types of “parasiteOf” or “parasitoidOf”, excluding social parasitism to focus on direct biological parasitism. Predation associations were extracted using interaction types of “preysOn” or “eats”. Given the relatively broad dietary range of predators, a genus-to-genus expansion was adopted for the predation relationships. Herbivorous interactions were extracted using the interaction types “hasHost” or “eats”, specifically targeting taxa within the kingdom Plantae. Extracted relationships were manually curated to verify the accuracy. Divergent times between species were extracted from TimeTree v5 via its application programming interface (API) (September 2023) (*56*). To test the effects of special features of sampled species pairs, three subsampling methods were employed using a custom Python script. For each method, 1000 replicates were randomly generated. For species pair shuffling, in every replication, the pairs of species were randomly rearranged to create new combinations. To control divergent time, pairs of species were randomly sampled from the background to match the divergent times in the tested category (i.e., parasitism, plant-sharing or predation). The sampling background pools were all species pairs excluding the specific tested category. A similar process was applied to control the last common ancestor. Linear model analyses, statistical tests, and data visualization were executed using R v4.3.1.

### Phylogenetic correction

We treated the minimum interspecific *wsp* distance between two species as a trait for each species pair (*i*, *j*). For any two pairs of species (*i*, *j*) and (*k*, *l*), we postulate that the covariance between their traits is given by: *Cov*[*Y_ij_,Y_kl_*]*=σ^2^*⋅(*T_ik_+T_jl_*),where *T_ik_*denotes the evolutionary time from the root to the most recent common ancestor (MRCA) of species *i* and *k*, and *Tjl* represents the evolutionary time from the root to the MRCA of species *j* and *l*. This formulation allows us to estimate the covariance structure of the traits, which is then integrated into our linear model analysis to adjust for phylogenetic non-independence. To manage computational constraints, we employed a random sampling strategy, selecting 1,000 species pairs from the “Others” category. We analyzed the relationship between the minimum interspecific *wsp* distance and group categories using a Generalized Least Squares model, accounting for the covariance structure. The model was implemented using the “gls” function in R package “nlme” v3.1.164.

### Ancestral state reconstruction

The taxonomic order of species from which wsp sequences originated was regarded as a discrete character trait. Ancestral state reconstruction was performed across the *wsp* phylogeny using the “ace” function in R package “phytools” v2.1.1. Three models were applied: Equal-Rates (ER), Symmetry (SYM), and All-Rates-Different (ARD). Among these three models, ARD was chosen as the best-fitting model, based on its lowest Akaike information criterion (AIC) value.

### Insect rearing

Both whiteflies *B. tabaci* and the parasitoid wasps *En. formosa* were maintained in nylon cages at 25±1°C, 70±5% RH, and a L:D photoperiod of 16:8 h. *Bemisia tabaci* was originally collected from the campus of Nanjing Agricultural University in 2012 and determined to be the Q biotype. An iso-female line of *B. tabaci*, which is naturally *Wolbachia*-free and has not been treated with antibiotics, was established. *En. formosa* was initially acquired from Beijing Ecoman Biotechnology Co., Ltd. in 2013 and was maintained on *B. tabaci* on tomato plants.

### PCR and Sanger sequencing

*Bemisia tabaci* or *En. formosa* were initially washed using ethanol and ultrapure water three times each. DNA was extracted from individuals using the STE method (*57*). To avoid false-negative results, a nested PCR targeting the *wsp* gene was employed for detecting *Wolbachia* in whiteflies (Fig. S8) as previously described (*57*). PCRs were performed using DreamTaq Green PCR Master Mix (Thermo Scientific) according to the manufacturer’s protocol. To confirm whether the *Wolbachia* present in whiteflies and their parasitoid wasps belonged to the same strain, PCR and Sanger sequencing were performed on *Wolbachia* genes, namely, *wsp*, *gatB*, *coxA*, *hcpA*, *ftsZ*, and *fbpA*, for MLST typing (*58*). The primers used for *wsp* nested PCR and MLST typing are listed in Table S1.

### FISH detection of *Wolbachia*

The location of *Wolbachia* in whiteflies was determined using fluorescence in situ hybridization as previously reported (*33*). To enhance the detection signal, two 5’ rhodamine-labeled probes, W1: 5′-AATCCGGCCGARCCGACCC-3′ and W2: 5′-CTTCTGTGAGTACCGTCATTATC-3′, were used to target *Wolbachia* 16S rRNA (*59*). Stained samples were examined under a Zeiss LSM 700 confocal microscope (Carl Zeiss, Germany). To confirm the specificity of the method, *Wolbachia*-free whiteflies that had not been parasitized by *En. formosa* were used as negative controls. These controls were processed in parallel with the *Wolbachia*-positive samples and imaged using identical exposure settings (Fig. S6).

### Outdoor cage experiments for *Wolbachia* transmission

To assess the potential transmission of *Wolbachia* from *En. formosa* to *B. tabaci*, six cages (80 cm x 80 cm x 80 cm, enclosed with a 120 mesh nylon screen) were established on the Nanjing Agricultural University campus. Initially, six 20-cm tall tomato plants were placed in each cage, with two additional plants introduced at 30-day intervals. Each cage was populated with 100 female and 100 male *Wolbachia*-free whiteflies. After a 20-day period, ten *En. formosa* wasps were introduced into each of three randomly selected cages, following the random sampling of 40 adult whiteflies from each cage. Subsequently, an additional 40 adult whiteflies were sampled from each cage at 20-day intervals. These sampled whiteflies were sexed based on their morphologies and screened for *Wolbachia* infection using *wsp* nested PCR.

### *Wolbachia* transmission from irradiated *En. formosa* to *B. tabaci*

To explore the possibility of *Wolbachia* transmission from *En. formosa* to *B. tabaci* due to unsuccessful parasitism, we subjected *En. formosa* wasps to radiation at the Nanjing Aerospace Irradiation Center. Wasps that had emerged within 24 h were exposed to ^60^Co radiation at doses of 60, 80 and 100 Gy (dose rate of 2 Gy/min). A single irradiated wasp was subsequently introduced into a Petri dish, which contained a tomato leaf infested with *Wolbachia*-free 3^rd^ or 4^th^ instar whitefly nymphs, and wet cotton was used to wrap the end of the leaf petiole to keep the leaf fresh. The parasitic behaviors of *En. formosa* were monitored under a Nikon SMZ800 stereomicroscope (Nikon Instrument Inc., Tokyo, Japan). Wasps were removed after 12 h of parasitism. Only whitefly nymphs that had been punctured by wasps were kept until the emergence of either parasitoid wasps or whiteflies. The newly emerged whiteflies were then collected for *Wolbachia* detection using *wsp* nested PCR. In each replication, five irradiated *En. formosa* were randomly selected for parasitization, with four replications performed in total. Nonirradiated wasps were used as controls.

### *Wolbachia* transmission across generations in whiteflies

The parasitized and subsequently emerged whiteflies were denoted as the initial generation (G_0_). To investigate the transmission rate of *Wolbachia* across whitefly generations, a pair of female and male newly emerged whitefly adults (G_0_) were randomly chosen and introduced into a Petri dish containing a tomato leaf. After 5 d of oviposition, the whitefly adults (G_0_) were removed and sampled for *Wolbachia* detection using *wsp* nested PCR. Only the eggs laid by *Wolbachia*-infected females were allowed to develop into adults. In a similar manner, upon the emergence of G_1_ adults, whiteflies were paired and placed into individual Petri dishes to produce the G_2_ generation and then sampled for *Wolbachia* detection. This procedure was repeated from the G_0_ to G_5_ generations. A parallel control experiment was conducted using *Wolbachia*-free whiteflies. Nine replications were conducted for each generation.

### Effects of *Wolbachia* on whitefly fitness

Given the inability to obtain a whitefly strain with 100% *Wolbachia* infection, we selected individuals from the infected whitefly population, which initially acquired *Wolbachia* through parasitism of irradiated *En. formosa* and subsequently maintained for over five generations. Whitefly nymphs were individually isolated into PCR tubes at their 4^th^ instar stage. After emergence, female and male whitefly adults were paired and allowed to oviposit for five days within a Petri dish containing a tomato leaf. These adults were then removed for *Wolbachia* detection via PCR. For the *Wolbachia*-infected female adults, we recorded several fitness parameters. These included the total number of eggs laid, the number of first instar nymphs, the developmental time from egg to eclosion, and the number of eclosed male and female offspring. This procedure was replicated with 60 pairs, and whiteflies from the original *Wolbachia*-free population were used as controls. Additionally, we examined the effects of various paternal and maternal infection status combinations on whitefly fitness. A similar procedure was employed, with the exception that all whitefly adults were sourced from the infected population, and their infection status was determined through *wsp* nested PCR.

### Statistics of the experimental data

Statistical analysis of experimental data was conducted using SPSS version 18.0 (SPSS Inc., Chicago, USA). The proportions were arcsine-square root transformed. Two independent samples were compared using Student’s T test. One-way ANOVA was performed for analysis of multiple samples. Means were then compared using Tukey’s honestly significant difference (HSD) test. Data were visualized using GraphPad Prism 6 software (GraphPad Software, San Diego, USA).

## Supporting information

source data for figures and tables

## Data and code availability

The alignment files, phylogenetic trees, and custom scripts can be accessed on FigShare (https://doi.org/10.6084/m9.figshare.24718119).

## Acknowledgements

This research was funded by “Shuangchuang Doctor” Foundation of Jiangsu Province (202030472), Nanjing Agricultural University startup fund (804018), the Hainan Major Science and Technology Project (ZDKJ2021007), and the Special Fund for Agro-scientific Research in the Public Interest of China (201303019). Bioinformatic analyses were supported by the high-performance computing platform of Bioinformatics Center, Nanjing Agricultural University.

## Author contributions

Y.X.L., X.Y.H., and Z.C.Y. conceptualized and designed the research; Z.C.Y. carried out the bioinformatics analysis; L.D.Q., H.L.J., and X.X.W. conducted the experiments; Y.X.L. and Z.C.Y. interpreted the results; Z.C.Y. and L.D.Q. prepared the visualization; Z.C.Y. wrote the initial manuscript; Y.X.L. and X.Y.H revised the manuscript.

## Supplementary Information

### Supplementary Note 1

We analyzed the relationship between the minimum interspecific *wsp* distance and group categories using a Generalized Least Squares model, accounting for the covariance structure. Given a dataset of 1,377 species, the formation of all possible species pairs results in an extensive 947,376 unique combinations. The construction of a comprehensive covariance matrix for these pairs would require the storage of an enormous 897,521,285,376 entries, which is beyond the capacity of standard computing systems. To reduce computational load, we randomly sampled 1,000 pairs from the total of 943,315 species pairs within the ‘Others’ category. After accounting for phylogenetic correction using covariance, both “Parasitism” and “Plant-sharing” had significant effects (both tests: *p* < 0.0001), while “Predation” showed no significant effect (*p* = 0.34). The estimated effects of “Parasitism” and “Plant-sharing” were -0.28 and -0.13, respectively.

### Supplementary Note 2

We divided all species pairs into six categories based on whether the two species belonged to the same genus, family, order, class, phylum, or kingdom. These categories represent different divergent levels between species pairs. Considering the impact of the divergent level, linear regression analyses were conducted with the *wsp* distance as the dependent variable and the category of divergent levels as the independent variable. We found that both “Parasitism” and “Plant-sharing” had significant effects (both tests: *p* < 2e^-16^), while “Predation” showed no significant effect (*p* = 0.58). The estimated effect of “Parasitism” (-0.28) was more profound than that of “Plant-sharing” (-0.14) (*p* = 1.32e^-15^).

### Supplementary Note 3

*Encarsia formosa* wasps that had emerged within 24 h were exposed to Co^60^ radiation at doses of 60, 80 and 100 Gy. On day 15 after irradiation treatments, the survival rate of adult wasps was 73.33 ± 3.33% in the nonirradiated group (Fig. S9). In contrast, the survival rates were 45.56 ± 1.92%, 7.78 ± 6.94%, and 3.33 ± 3.33% after exposure to 60 Gy, 80 Gy, and 100 Gy radiation, respectively. The survival rate of adult wasps decreased with increasing irradiation doses (Fig. S9; ANOVA; *F_3,8_* = 34.18, *p* < 0.001). After irradiation, the ovaries were dissected, and changes in egg dimensions were recorded (Fig. S10). Following 80 and 100 Gy irradiation, almost no eggs were detected starting from day 4, while after 60 Gy irradiation, eggs were nearly absent starting from day 9. Fig. S11 shows the ovaries on days 1, 4, and 9 post-exposure to various irradiation doses. In the nonirradiated *En. formosa*, there were no obvious changes in the morphology of the ovarioles or oocytes from days 1 to 9 (Fig. S11). However, 4 days post-irradiation, the ovarioles of wasps exposed to 60 Gy appeared thinner, and some oocytes had dissolved (Fig. S11). In addition, the ovarioles of wasps exposed to 80 Gy and 100 Gy had become transparent, with the majority of the oocytes disappearing and becoming invisible (Fig. S11). Parasitism experiments demonstrated that the nonirradiated *En. formosa* laid 7.3 ± 1.1 eggs in 12 h. This number significantly dropped to 3.2 ± 0.9 eggs for wasps exposed to 60 Gy radiation (Fig. 3a; Student’s t test; *t* = 12.91, *p* < 0.001). The wasps exposed to 80 Gy and 100 Gy radiation almost completely ceased egg laying, resulting in insufficient data collection.

---

**Table S1.**
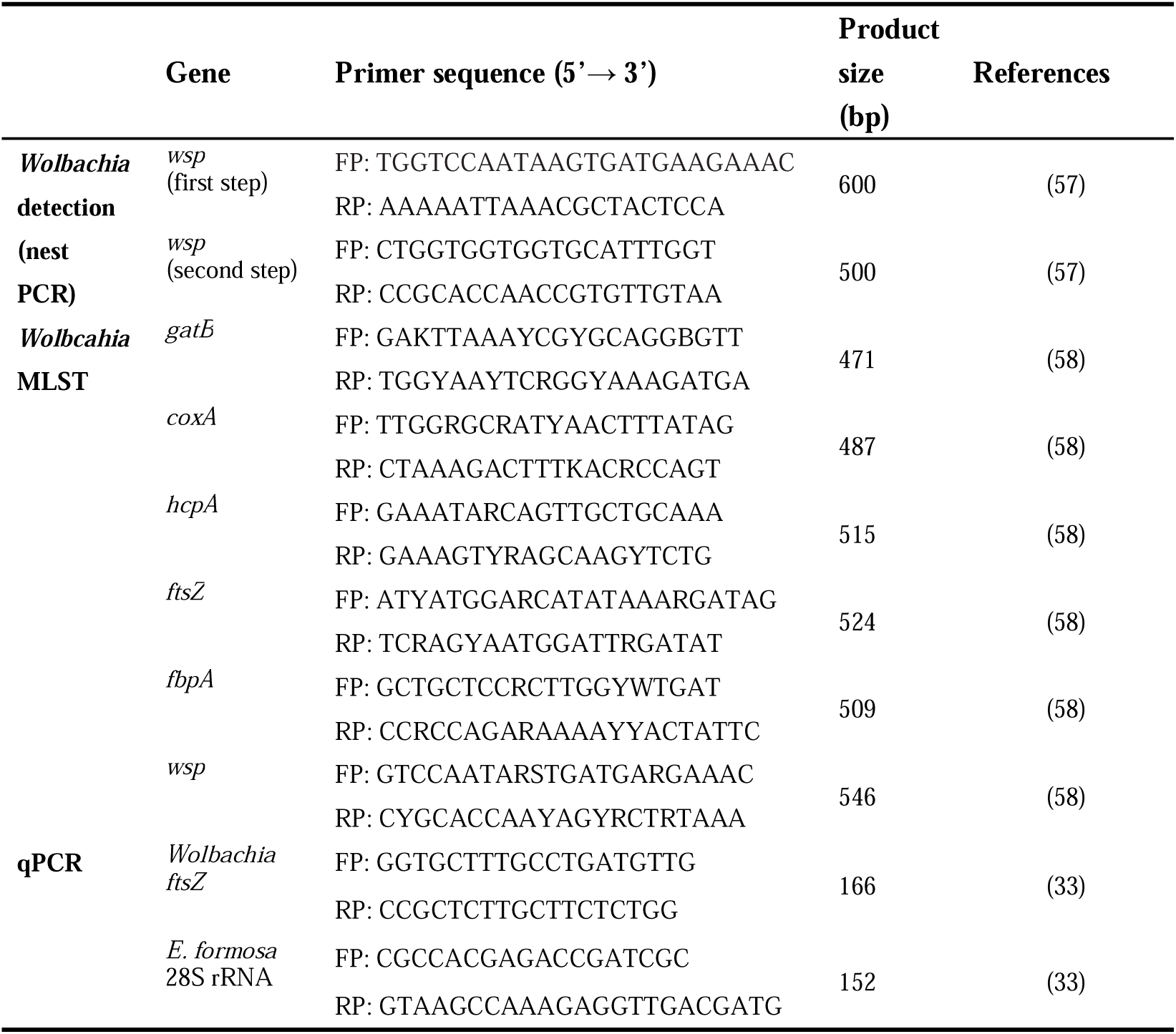
Primers used in this study.

**Fig. S1.**
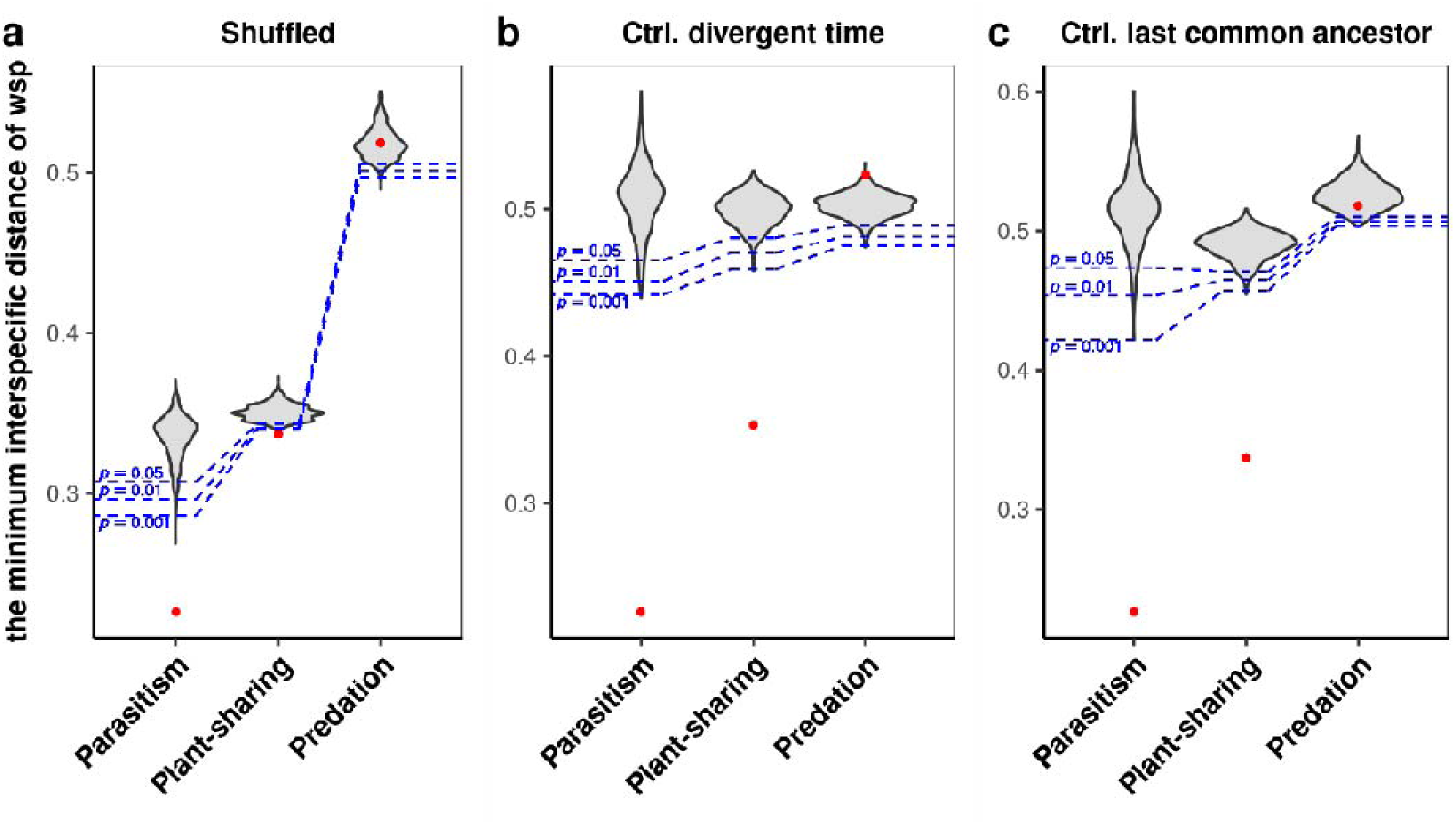
Subsampling by (a) shuffling species pairs, (b) controlling divergent time or (c) last common ancestor. For each method, 1000 replicates were randomly generated. Details are provided in the Methods and Materials. Red dots indicate the observed minimum interspecific distances of *wsp* for different species pair categories.

**Fig. S2.**
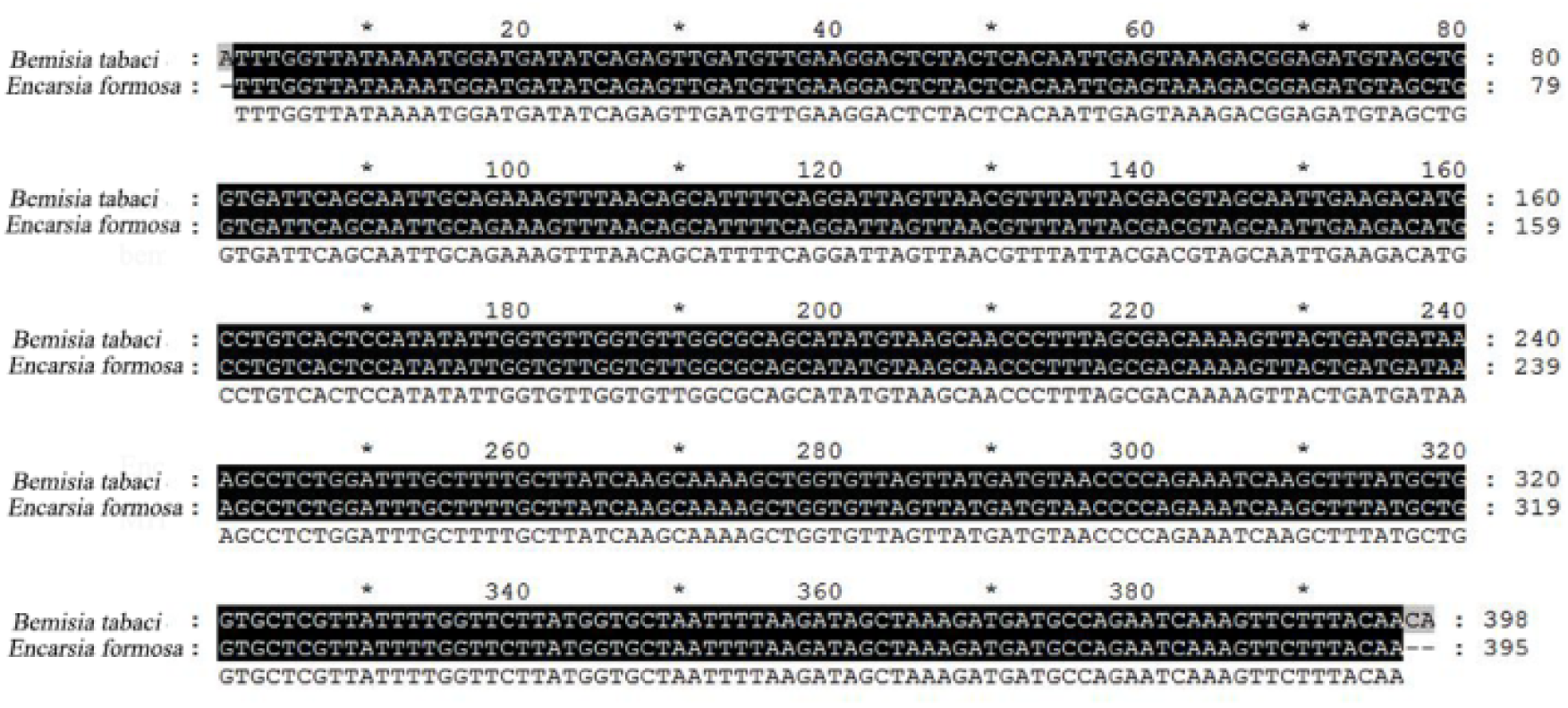
Alignment of wsp sequences detected from *En. formosa* and *B. tabaci* in the cage experiment.

**Fig. S3.**
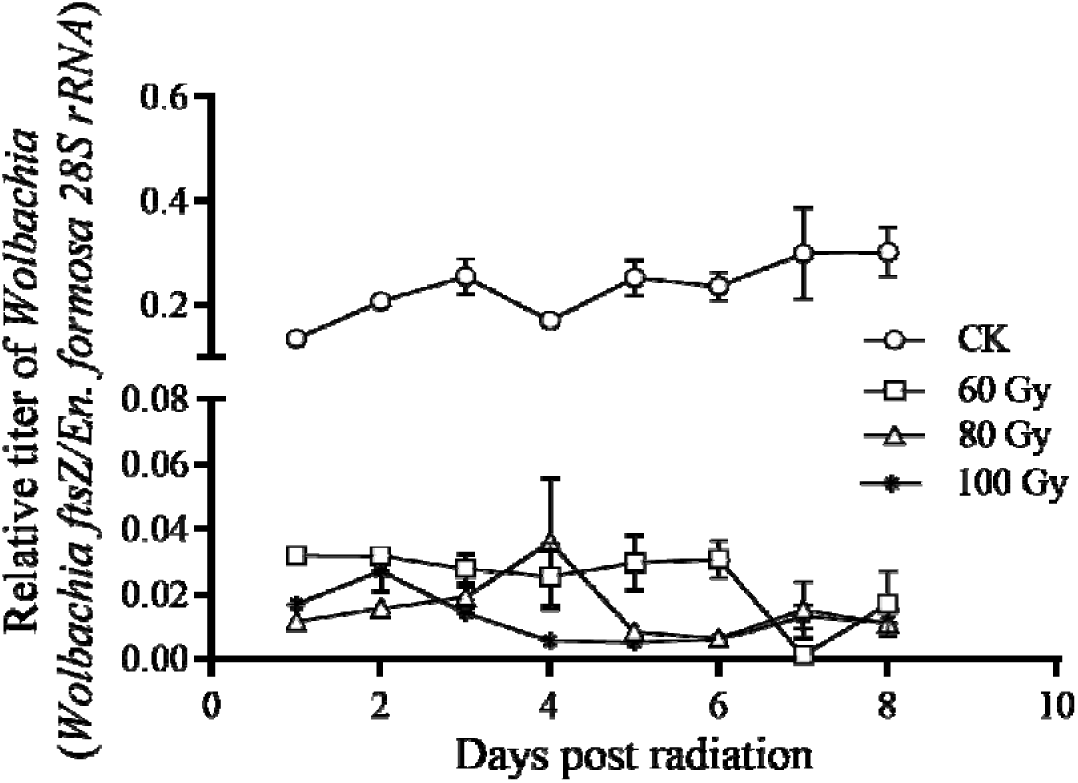
Relative titer of *Wolbachia* in *En. formosa* after radiation. Relative *Wolbachia* titer in radiated female adults of *En. formosa* were evaluated using qPCR of the *Wolbachia ftsZ* gene. The 28S rRNA gene of *En. formosa* was used as the reference. Data are presented as the means ± SEs (n = 5).

**Fig. S4.**
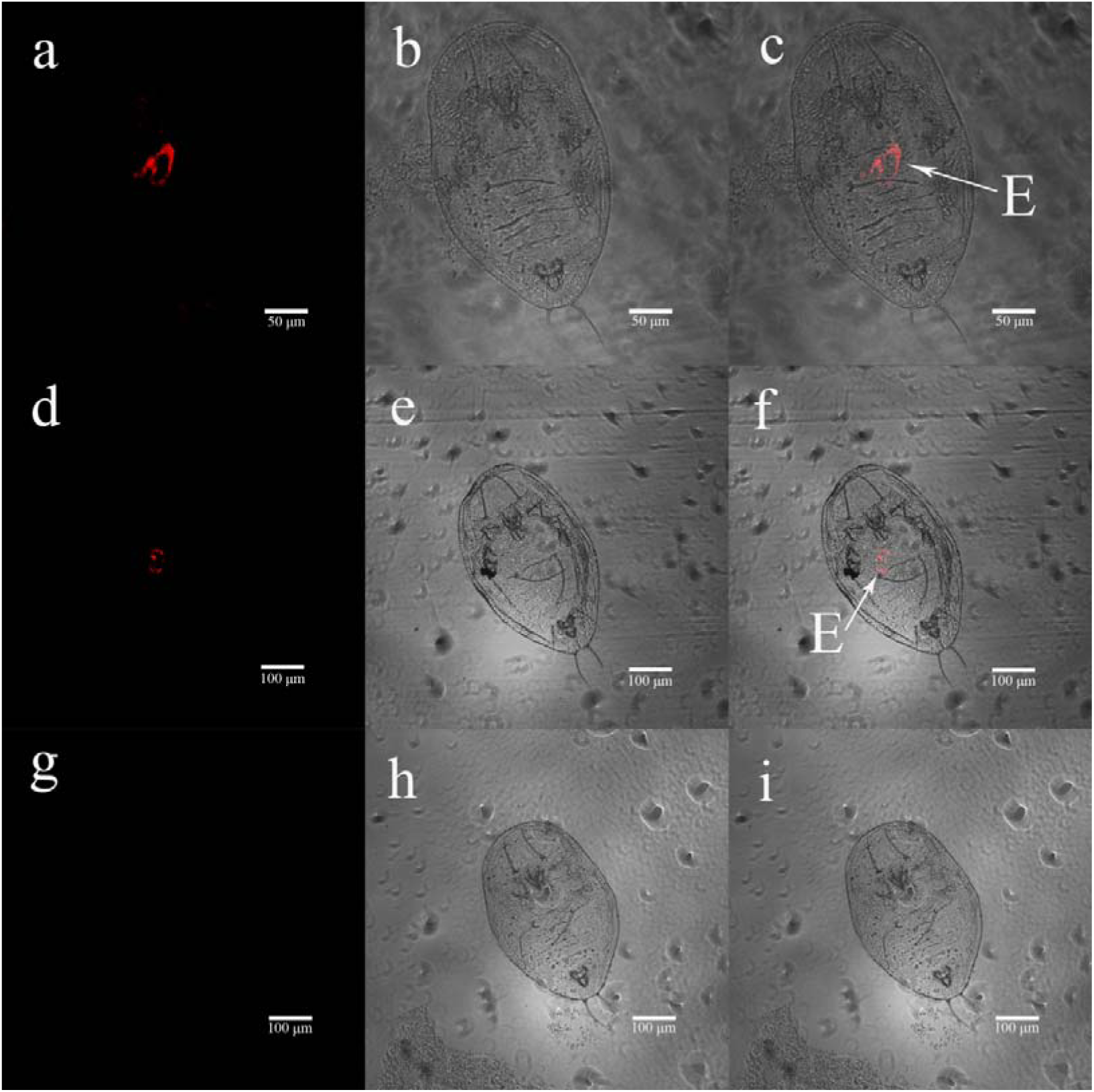
FISH visualization of *Wolbachia* in parasitized and nonparasitized nymphs of *B. tabaci*. **(a–c)** Fluorescence **(a)**, bright field **(b)** and combined **(c)** images of parasitized 2nd instar nymph of *B. tabaci*. **(d–f)** Fluorescence **(d)**, bright field **(e)** and combined **(f)** images of parasitized 3rd instar nymph of *B. tabaci*. **(g–i)** Fluorescence **(g)**, bright field **(h)** and combined **(i)** images of nonparasitized 2nd instar nymphs of *B. tabaci*. The merged images in panels c, f, and i correspond to Fig. 3f, e, and d, respectively. E: injected eggs from *En. formosa*.

**Fig. S5.**
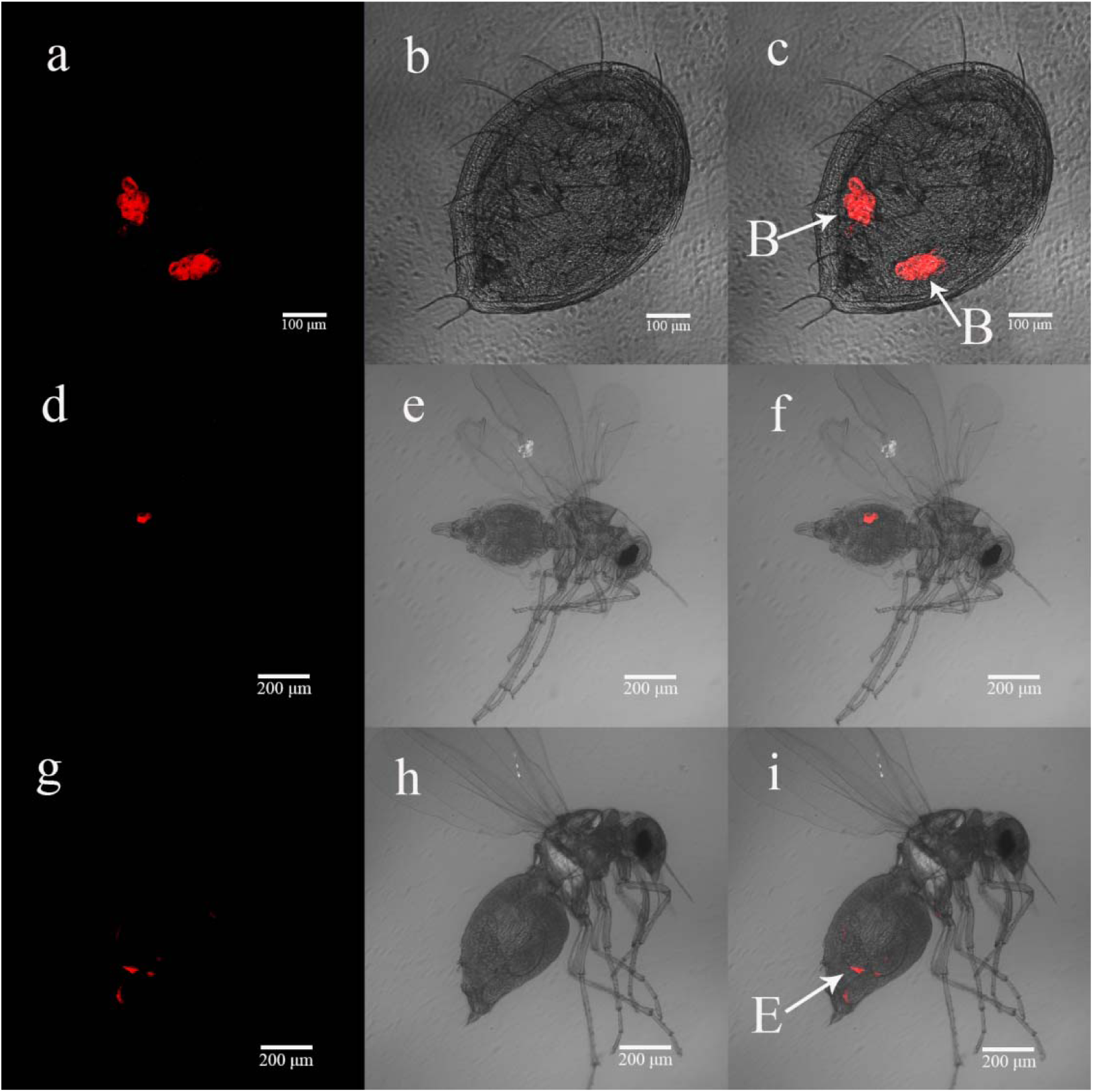
FISH visualization of *Wolbachia* in G_3_ *B. tabaci*. **(a–c)** Fluorescence **(a)**, bright field **(b)** and combined **(c)** of G_3_ whitefly nymph. **(d–f)** Fluorescence **(d)**, bright field **(e)** and combined **(f)** images of G_3_ whitefly male adults. **(g–i)** Fluorescence **(g)**, bright field **(h)** and combined **(i)** images of G_3_ whitefly female adults. The merged images in panels c, f, and i correspond to Fig. 4c, d, and e, respectively. B: structures morphologically resembling bacteriocytes; E: eggs in ovary of female whitefly.

**Fig. S6.**
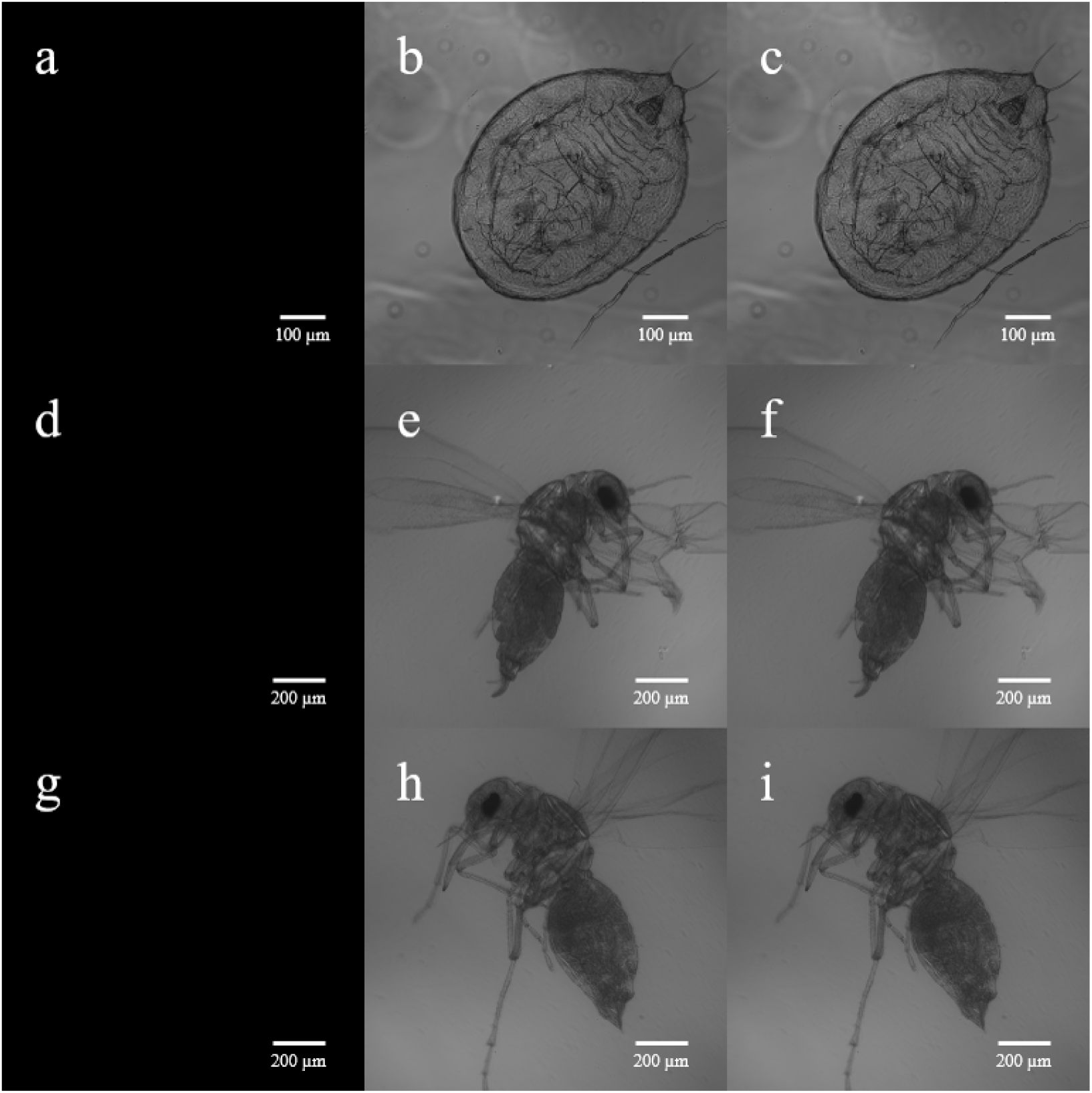
FISH analysis of nonparasitized, *Wolbachia*-free *B. tabaci* as negative controls. **(a–c)** Fluorescence **(a)**, bright field **(b)** and combined **(c)** images of a whitefly nymph. **(d–f)** Fluorescence **(d)**, bright field **(e)** and combined **(f)** images of a whitefly male adult. **(g–i)** Fluorescence **(g)**, bright field **(h)** and combined **(i)** images of a whitefly female adult. All negative-control whiteflies were *Wolbachia*-free and had not been parasitized by *En. formosa*. They were processed in parallel with *Wolbachia*-positive samples and imaged using identical exposure settings. No *Wolbachia*-specific FISH signals were detected. Scale bars: 100 µm (a–c) and 200 µm (d–i).

**Fig. S7.**
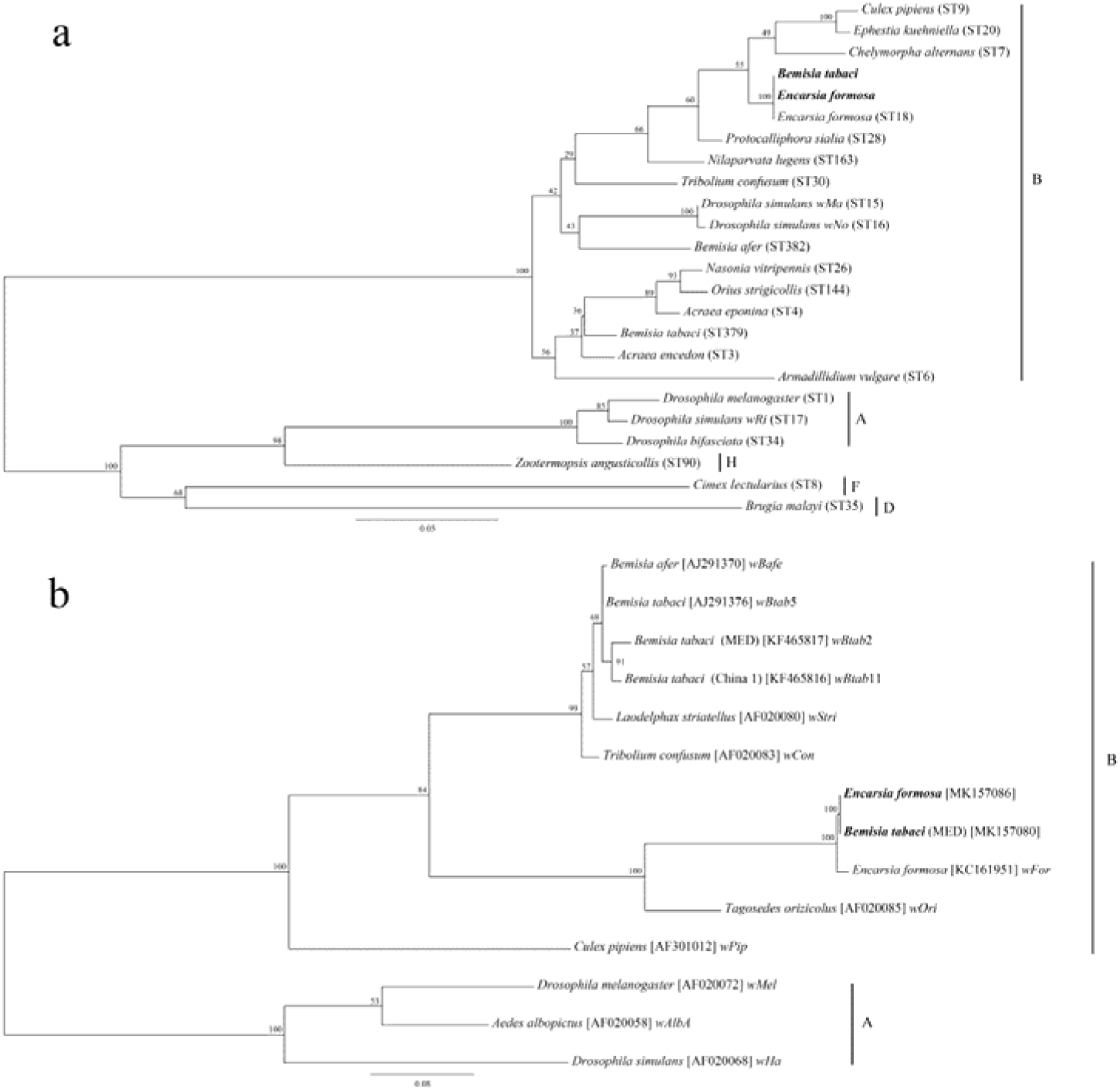
Maximum likelihood phylogenetic analysis of detected *Wolbachia* strains in *B. tabaci* and *En. formosa*. **(a)** Phylogenetic tree constructed using five MLST genes (i.e., *coxA*, *fbpA*, *gatB*, *hcpA*, *ftsZ*). **(b)** Phylogenetic tree constructed using the *wsp* gene. The bootstrap values are represented at the nodes. Sequences obtained in this study are emphasized in bold.

**Fig. S8.**
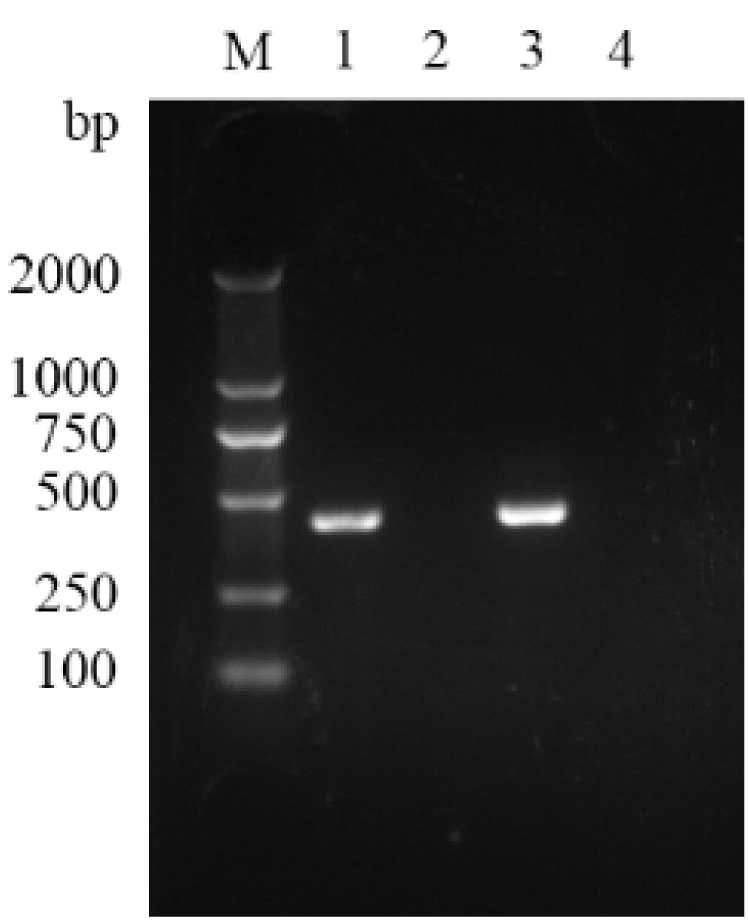
PCR detection of *Wolbachia* in *B. tabaci*. M: DNA marker; lane 1: *Wolbachia*-positive whitefly; lane 2: *Wolbachia*-negative whitefly; lane 3: *En. formosa* as a positive control; lane 4: ddH_2_O as a negative control.

**Fig. S9.**
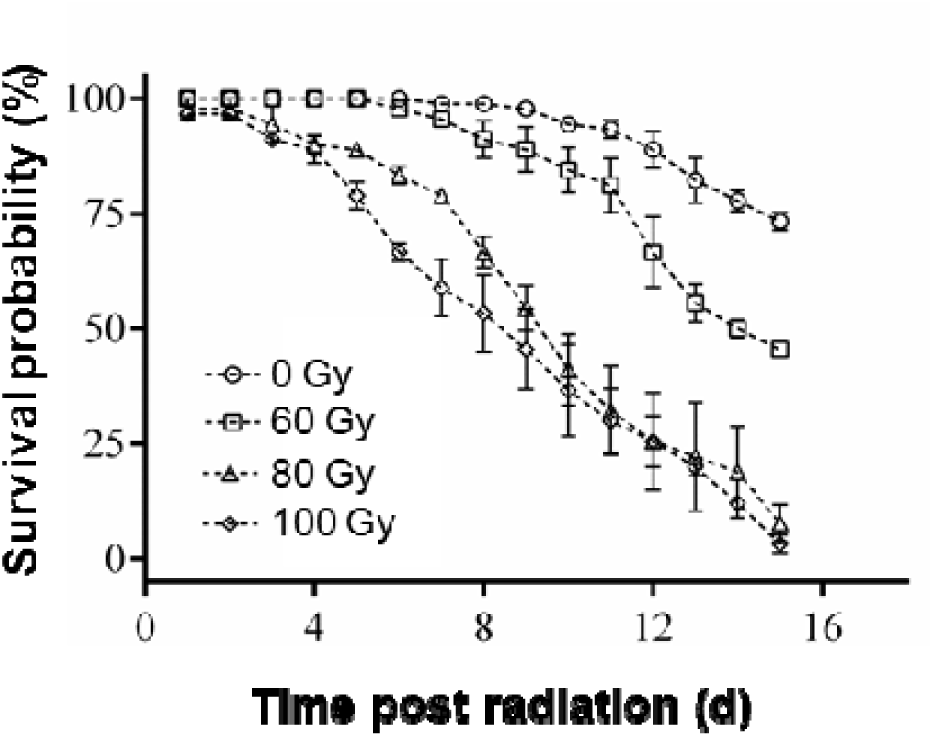
Survival of *En. formosa* after radiation at various doses. Data are presented as the means ± SEs (n = 8).

**Fig. S10.**
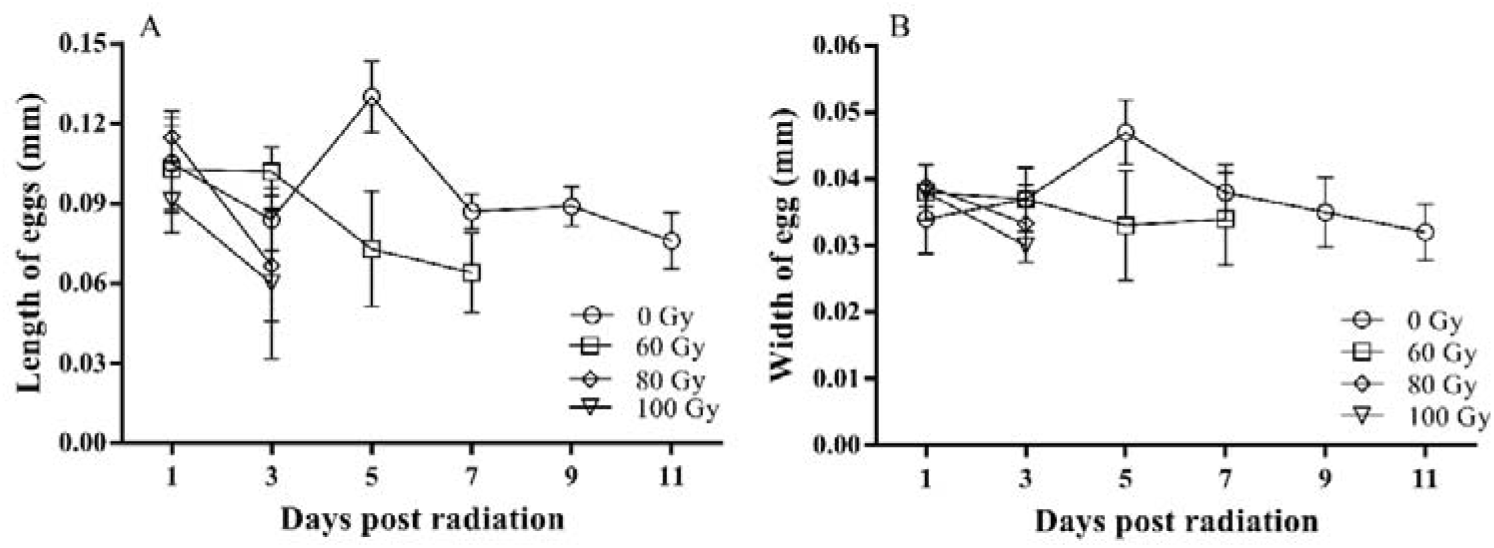
Lengths (a) and widths (b) of *En. formosa* eggs after irradiation at various doses. Data are presented as the means ± SEs. The sample sizes are typically 10, with the exception of day 3 post-irradiation at 80 Gy and 100 Gy, where the sample sizes are 3 and 2, respectively. This reduction in sample size is due to the scarcity of eggs resulting from irradiation.

**Fig. S11.**
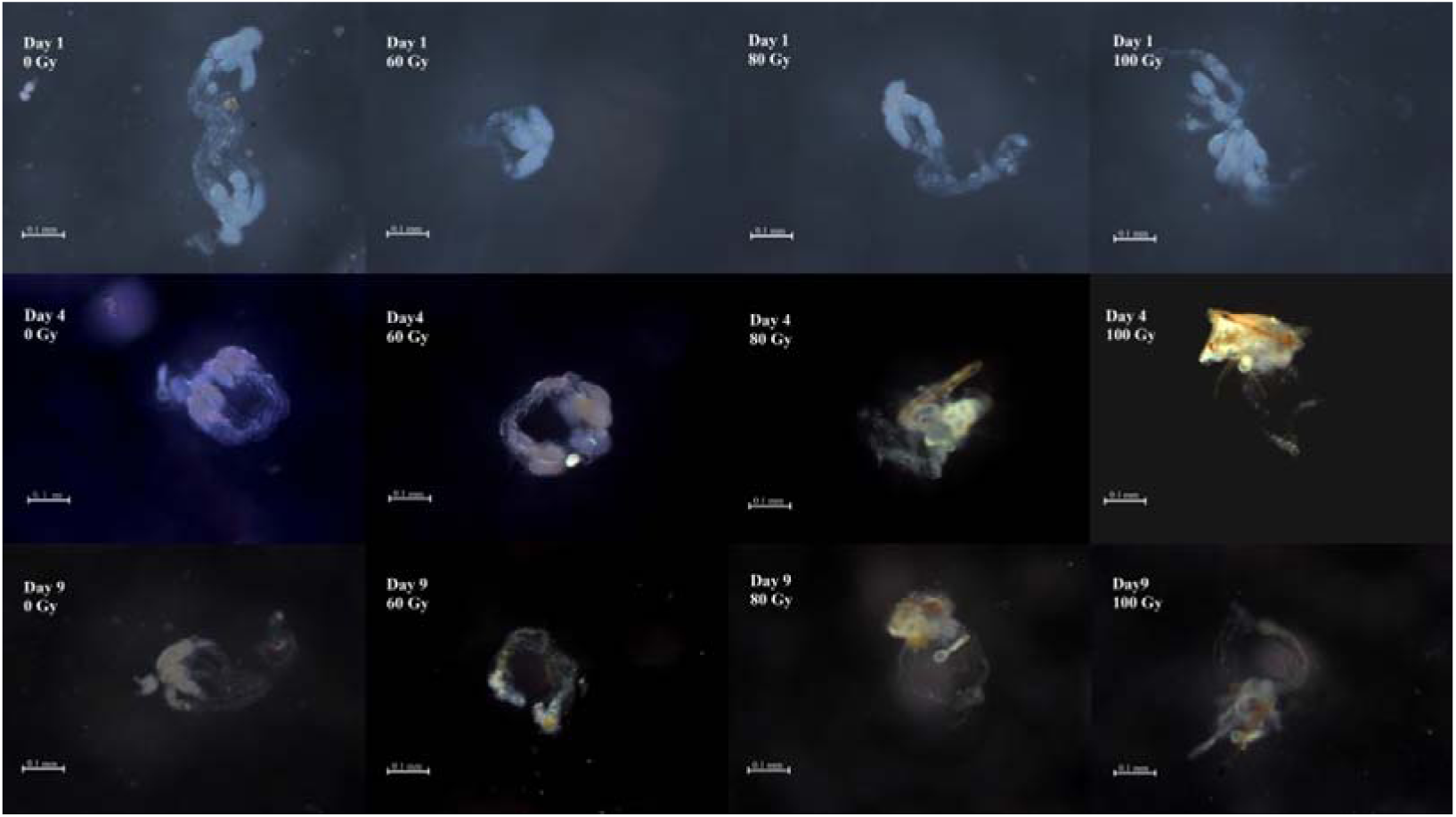
Morphologies of dissected *En. formosa* ovaries at 1, 4 and 9 days after radiation at various doses.

## References

1. M. Bright, S. Bulgheresi, A complex journey: transmission of microbial symbionts. Nat Rev Microbiol 8, 218–230 (2010).

2. J. Perreau, N. A. Moran, Genetic innovations in animal-microbe symbioses. Nat Rev Genet 23, 23–39 (2022).

3. A. E. Douglas, Multiorganismal insects: diversity and function of resident microorganisms. Annu Rev Entomol 60, 17–34 (2015).

4. A. Mestre, R. Poulin, J. Hortal, A niche perspective on the range expansion of symbionts. Biol Rev Camb Philos Soc 95, 491–516 (2020).

5. J. L. Sachs, R. G. Skophammer, J. U. Regus, Evolutionary transitions in bacterial symbiosis. Proc Natl Acad Sci USA 108 **Suppl 2**, 10800–10807 (2011).

6. R. M. Fisher, L. M. Henry, C. K. Cornwallis, E. T. Kiers, S. A. West, The evolution of host-symbiont dependence. Nat Commun 8, 15973 (2017).

7. J. H. Werren, L. Baldo, M. E. Clark, *Wolbachia:* master manipulators of invertebrate biology. Nat Rev Microbiol 6, 741–751 (2008).

8. R. Kaur et al., Living in the endosymbiotic world of *Wolbachia*: a centennial review. Cell Host Microbe 29, 879–893 (2021).

9. J. Porter, W. Sullivan, The cellular lives of *Wolbachia*. Nat Rev Microbiol 21, 750–766 (2023).

10. M. Scholz et al., Large scale genome reconstructions illuminate *Wolbachia evolution*. Nat Commun 11, 5235 (2020).

11. E. Vancaester, M. Blaxter, Phylogenomic analysis of Wolbachia genomes from the Darwin Tree of Life biodiversity genomics project. PLoS Biol 21, e3001972 (2023).

12. X. Zheng et al., Incompatible and sterile insect techniques combined eliminate mosquitoes. Nature 572, 56–61 (2019).

13. A. A. Hoffmann et al., Successful establishment of Wolbachia in Aedes populations to suppress dengue transmission. Nature 476, 454–457 (2011).

14. J. T. Gong, T. P. Li, M. K. Wang, X. Y. Hong, Wolbachia-based strategies for control of agricultural pests. Curr Opin Insect Sci 57, 101039 (2023).

15. J. T. Gong et al., Stable Introduction of Plant-Virus-Inhibiting Wolbachia into Planthoppers for Rice Protection. Curr Biol 30, 4837–4845.e4835 (2020).

16. E. L. Loreto, G. L. Wallau, Risks of Wolbachia mosquito control. Science 351, 1273 (2016).

17. E. Sanaei, S. Charlat, J. Engelstadter, *Wolbachia* host shifts: routes, mechanisms, constraints and evolutionary consequences. Biol Rev Camb Philos Soc 96, 433–453 (2021).

18. H. Noda et al., Infection shared among planthoppers (Homoptera: Delphacidae) and their endoparasite (Strepsiptera: Elenchidae): a probable case of interspecies transmission. Mol Ecol 10, 2101–2106 (2001).

19. F. Vavre, F. Fleury, D. Lepetit, P. Fouillet, M. Bouletreau, Phylogenetic evidence for horizontal transmission of Wolbachia in host-parasitoid associations. Mol Biol Evol 16, 1711–1723 (1999).

20. P. Kittayapong, W. Jamnongluk, A. Thipaksorn, J. R. Milne, C. Sindhusake, Infection complexity among insects in the tropical rice-field community. Mol Ecol 12, 1049–1060 (2003).

21. H. Hou, G. Zhao, C. Su, D. Zhu, Wolbachia prevalence patterns: horizontal transmission, recombination, and multiple infections in chestnut gall wasp-parasitoid communities. Entomol Exp Appl 168, 752–765 (2020).

22. J. L. Morrow, M. Frommer, D. C. A. Shearman, M. Riegler, Tropical tephritid fruit fly community with high incidence of shared strains as platform for horizontal transmission of endosymbionts. Environ Microbiol 16, 3622–3637 (2014).

23. D. Kageyama, S. Narita, T. Imamura, A. Miyanoshita, Detection and identification of *Wolbachia* endosymbionts from laboratory stocks of stored-product insect pests and their parasitoids. J Stored Prod Res 46, 13–19 (2010).

24. B. D. Heath, R. D. J. Butcher, W. G. F. Whitfield, S. F. Hubbard, Horizontal transfer of *Wolbachia* between phylogenetically distant insect species by a naturally occurring mechanism. Curr Biol 9, 313–316 (1999).

25. M. Aboumourad, H. zu Dohna, Unidirectional and heterogenous Wolbachia transfer rates among insect host orders. Preprint at https://www.researchsquare.com/article/rs-2698051/v1, (2023).

26. D. P. Hughes, P. Pamilo, J. Kathirithamby, Horizontal transmission of Wolbachia by strepsipteran endoparasites? A response to Noda et al., 2001. Mol Ecol 13, 507–509 (2004).

27. M. Z. Ahmed et al., The intracellular bacterium Wolbachia uses parasitoid wasps as phoretic vectors for efficient horizontal transmission. PLoS Pathog 11, e1004672 (2015).

28. M. P. Hassell, J. K. Waage, Host-parasitoid population interactions. Annu Rev Entomol 29, 89–114 (1984).

29. K. M. Oliver, J. A. Russell, N. A. Moran, M. S. Hunter, Facultative bacterial symbionts in aphids confer resistance to parasitic wasps. Proc Natl Acad Sci USA 100, 1803–1807 (2003).

30. S. M. Smith, Biological control with *Trichogramma:* advances, successes, and potential of their use. Annu Rev Entomol 41, 375–406 (1996).

31. L. Qi, J. Sun, X. Hong, Y. Li, Diversity and Phylogenetic Analyses Reveal Horizontal Transmission of Endosymbionts Between Whiteflies and Their Parasitoids. J Econ Entomol 112, 894–905 (2019).

32. M. S. Hoddle, R. G. Van Driesche, J. P. Sanderson, Biology and use of the whitefly parasitoid *Encarsia formosa*. Annu Rev Entomol 43, 645–669 (1998).

33. X. X. Wang, L. D. Qi, R. Jiang, Y. Z. Du, Y. X. Li, Incomplete removal of Wolbachia with tetracycline has two-edged reproductive effects in the thelytokous wasp *Encarsia formosa* (Hymenoptera: Aphelinidae). Sci Rep 7, 44014 (2017).

34. S. A. Andreason et al., Whitefly Endosymbionts: Biology, Evolution, and Plant Virus Interactions. Insects 11, 775 (2020).

35. X. L. Bing et al., Diversity and evolution of the *Wolbachia* endosymbionts of Bemisia (Hemiptera: Aleyrodidae) whiteflies. Ecol Evol 4, 2714–2737 (2014).

36. E. Zchori-Fein, T. Lahav, S. Freilich, Variations in the identity and complexity of endosymbiont combinations in whitefly hosts. Front Microbiol 5, 310 (2014).

37. G. Gueguen et al., Endosymbiont metacommunities, mtDNA diversity and the evolution of the *Bemisia tabaci* (Hemiptera: Aleyrodidae) species complex. Mol Ecol 19, 4365–4376 (2010).

38. M. Z. Ahmed, J. W. Breinholt, A. Y. Kawahara, Evidence for common horizontal transmission of *Wolbachia* among butterflies and moths. BMC Evol Biol 16, 118 (2016).

39. M. E. Huigens et al., Infectious parthenogenesis. Nature 405, 178–179 (2000).

40. S. J. Li et al., Plant-mediated horizontal transmission of Wolbachia between whiteflies. ISME J 11, 1019–1028 (2017).

41. J. Popovici et al., Assessing key safety concerns of a *Wolbachia*-based strategy to control dengue transmission by *Aedes mosquitoes*. Mem. Inst. Oswaldo Cruz 105, 957–964 (2010).

42. V. G. Faria, T. F. Paulo, É. Sucena, Testing cannibalism as a mechanism for horizontal transmission of *Wolbachia* in *Drosophila*. Symbiosis 68, 79–85 (2016).

43. Q. Su, G. Hu, Y. Yun, Y. Peng, Horizontal transmission of *Wolbachia* in *Hylyphantes graminicola* is more likely via intraspecies than interspecies transfer. Symbiosis 79, 123–128 (2019).

44. E. Chiel et al., Characteristics, phenotype, and transmission of Wolbachia in the sweet potato whitefly, Bemisia tabaci (Hemiptera: Aleyrodidae), and its parasitoid Eretmocerus sp. nr. emiratus (Hymenoptera: Aphelinidae). Environ Entomol 43, 353–362 (2014).

45. I. D. Weber, C. Czepak, K. C. Albernaz-Godinho, K. M. B. Borges, A. S. G. Coelho, Validation of a sampling method: using one square centimeter for sampling the immature stages of Bemisia tabaci in soybean. Entomol Exp Appl 170, 488–494 (2022).

46. P. Shi et al., *Wolbachia* has two different localization patterns in whitefly *Bemisia tabaci* AsiaII7 Species. PLoS One 11, e0162558 (2016).

47. R. Arunkumar et al., Natural selection has driven the recurrent loss of an immunity gene that protects *Drosophila* against a major natural parasite. Proc Natl Acad Sci USA 120, e2211019120 (2023).

48. J. J. Zhang et al., Effects of host-egg ages on host selection and suitability of four Chinese Trichogramma species, egg parasitoids of the rice striped stem borer, Chilo suppressalis. Biocontrol 59, 159–166 (2014).

49. M. E. Huigens, R. P. de Almeida, P. A. Boons, R. F. Luck, R. Stouthamer, Natural interspecific and intraspecific horizontal transfer of parthenogenesis-inducing *Wolbachia* in *Trichogramma* wasps. Proc Biol Sci 271, 509–515 (2004).

50. K. Katoh, D. M. Standley, MAFFT multiple sequence alignment software version 7: improvements in performance and usability. Mol Biol Evol 30, 772–780 (2013).

51. M. Suyama, D. Torrents, P. Bork, PAL2NAL: robust conversion of protein sequence alignments into the corresponding codon alignments. Nucleic Acids Res 34, W609–612 (2006).

52. B. Q. Minh et al., IQ-TREE 2: new models and efficient methods for phylogenetic inference in the genomic era. Mol Biol Evol 37, 1530–1534 (2020).

53. S. Kalyaanamoorthy, B. Q. Minh, T. K. F. Wong, A. von Haeseler, L. S. Jermiin, ModelFinder: fast model selection for accurate phylogenetic estimates. Nat Methods 14, 587–589 (2017).

54. I. Letunic, P. Bork, Interactive tree of life (iTOL) v3: an online tool for the display and annotation of phylogenetic and other trees. Nucleic Acids Res 44, W242–245 (2016).

55. J. H. Poelen, J. D. Simons, C. J. Mungall, Global biotic interactions: An open infrastructure to share and analyze species-interaction datasets. Ecol. Inform. 24, 148–159 (2014).

56. S. Kumar et al., TimeTree 5: an expanded resource for species divergence times. Mol Biol Evol 39, msac174 (2022).

57. H. L. Ji, L. D. Qi, X. Y. Hong, H. F. Xie, Y. X. Li, Effects of host sex, plant species, and putative host species on the prevalence of *Wolbachia* in natural populations of *Bemisia tabaci* (Hemiptera: Aleyrodidae): a modified nested PCR study. J Econ Entomol 108, 210–218 (2015).

58. L. Baldo et al., Multilocus sequence typing system for the endosymbiont *Wolbachia pipientis*. Appl Environ Microbiol 72, 7098–7110 (2006).

59. A. Heddi, A. M. Grenier, C. Khatchadourian, H. Charles, P. Nardon, Four intracellular genomes direct weevil biology: nuclear, mitochondrial, principal endosymbiont, and *Wolbachia*. Proc Natl Acad Sci USA 96, 6814–6819 (1999).

